# Invasion preferences suggest a possible role for *Plasmodium falciparum* parasites in the expansion of Duffy negativity in West and Central Africa

**DOI:** 10.1101/2025.07.09.663497

**Authors:** Balanding Manneh, Viola Introini, James Reed, Madalina Rotariu, Robin Antrobus, Pietro Cicuta, Michael P. Weekes, Bridget S. Penman, Julian C. Rayner

**Author notes:** Correspondence (Julian C. Rayner).

## Abstract

Duffy antigen receptor for chemokines (DARC) is the primary red blood cell (RBC) receptor for invasion of human RBCs by *Plasmodium vivax* and *knowlesi* parasites. By contrast, *Plasmodium falciparum* parasites use multiple RBC receptors for invasion. Whether DARC is one of these receptors has never been systematically explored. We used flow cytometry and microscopy-based approaches to investigate whether *P. falciparum* parasites preferentially invade specific Duffy RBC phenotypes and explored two potential explanations for invasion preference – differences in RBC biophysical properties and surface protein composition. *P. falciparum* parasites showed a consistent preference for Duffy-positive RBCs, and some biophysical properties and surface protein expression varied between Duffy-positive and Duffy-negative RBCs. We then used our *in vitro* invasion data to parametrise an evolutionary-epidemiological model of the relationship between *P. falciparum* and the FYB^ES^ allele. Our model accounts for immunity against *P. falciparum* virulence, gained through exposure, and thus mutations that impede infection are not always advantageous. The inhibition of *P. falciparum* invasion that we observed *in vitro* leads to FYB^ES^ frequencies increasing at low levels of *P. falciparum* transmission but decreasing at high levels of transmission. The impact of *P. falciparum* on the prevalence of Duffy negativity may therefore be most apparent in lower transmission settings. Our findings are the first to show a link between Duffy negativity and *P. falciparum* and suggest that DARC may direct or indirectly be involved in *P. falciparum* invasion of human RBCs which could, together with *P. vivax*, explain the distribution of Duffy negativity in sub-Saharan Africa.

## INTRODUCTION

Humans are genetically diverse, and this diversity can impact their susceptibility to infectious diseases (Allison 1954; Roberts and Williams 2013). Malaria in particular has acted as a strong force shaping the human genome and has selected for multiple natural variants which provide varying degrees of protection against malaria and are therefore found at higher frequencies in malaria-endemic regions (Williams 2006). The beta globin mutation responsible for sickle haemoglobin (HbS), found at a minor allele frequency (MAF) of up to 18% in some parts of sub-Saharan Africa, is perhaps the most well-studied – it causes sickle cell disease in homozygotes (HbSS) but confers significant protection against malaria in heterozygotes (HbAS)(Allison 1954; Piel et al. 2010; Grosse et al. 2011). However, numerous other polymorphisms that alter RBC structure and function have also been associated with protection against severe malaria; they include both relatively common (e.g., O Blood Group and Glucose-6-Phosphate Deficiency) and rare variants (e.g., Dantu blood group and variants in the RBC calcium transporter, ATP2B4). Such variants have been shown to mediate protection via a range of mechanisms including reducing parasite invasion of RBCs, slowing parasite growth after invasion has occurred, and/or reducing the ability of infected RBCs to cytoadhere to the microvasculature (Rowe et al. 2007; Manjurano et al. 2015; Lessard et al. 2017; Zámbó et al. 2017; Kariuki et al. 2020; Kariuki et al. 2023).

While the majority of studies on malaria resistance and susceptibility have focused on *Plasmodium falciparum* malaria, the Duffy blood group system is an example that has long been strongly associated with protection against *Plasmodium vivax* malaria (Miller et al. 1976). The Duffy blood group antigens are contained within the Duffy antigen receptor for chemokines (DARC), a seven-transmembrane glycoprotein that is expressed on the surfaces of many cell types, including RBCs and endothelial cells, and is encoded by the Duffy (Fy) gene on chromosome 1 (Donahue et al. 1968; Peiper et al. 1995). The Duffy blood group consists of the high-frequency FYA, FYB, and FYB^ES^ alleles (*ES = erythrocyte silent*) that give rise to four major phenotypes: Fy(a+b-), Fy(a-b+), Fy(a+b+), and Fy(a-b-) as well as multiple other rare alleles (International Society of Blood Transfusion (ISBT) 2021). The co-dominant FYA and FYB alleles encode the Fy^a^ and Fy^b^ Duffy blood group antigens, which differ due to a single missense mutation (125G>A; rs12075) and result in an amino acid change at residue 42. Glycine at this residue defines Fy^a^, while Aspartic Acid defines the Fy^b^ antigen (Langhi and Bordin 2006). FYB^ES^ is defined by a single nucleotide substitution (-67T>C, rs2814778; previously -33T>C) in the 5’UTR/promoter region, within a recognition motif for GATA1 (erythroid transcription factor) in the ancestral FYB allele. This substitution was thought to completely abolish the expression of DARC on RBCs but not on other cell types (Chaudhuri et al. 1995; Höher et al. 2018); however, a recent study reported DARC expression in the erythroid precursors of genotypically Duffy-negative individuals (Dechavanne et al. 2023). FYB^ES^ has a frequency of >90% in parts of West and Central Africa, and homozygosity for the allele produces the Duffy-negative RBC phenotype, Fy(a-b-)(Howes et al. 2011; Höher et al. 2018).

The absence of the DARC receptor on RBCs of Duffy-negative individuals makes them largely resistant to *P. vivax* infection, as *P. vivax* relies almost exclusively on this receptor for RBC invasion (Miller et al. 1976). This reliance was for many years thought to be absolute, but recent studies conducted in Madagascar (Ménard et al. 2010), Mauritania (Wurtz et al. 2011), Brazil (Carvalho et al. 2012) and elsewhere (Twohig et al. 2019) have all reported the presence of *P. vivax* in Duffy-negative individuals, suggesting that either *P. vivax* is evolving to invade RBCs of Duffy-negative individuals using an alternate receptor (Lee et al. 2024), or that such infections have always been historically present but not previously detected.

Previous studies on the links between DARC and malaria have therefore predominantly focused on its role as a receptor for *P. vivax* and the related zoonotic pathogen *P. knowlesi*, where the interaction between DARC and its cognate parasite binding partners (Duffy Binding Protein (DBP); *Pv*DBP and *Pk*DBP), plays a dominant role during human RBC invasion (Batchelor et al. 2014). In contrast, *P. falciparum* is well known to invade the RBCs of both Duffy-positive and Duffy-negative individuals and relies on multiple RBC receptors, including the glycophorins (GYPA, GYPB, and GYPC), to facilitate the RBC invasion process (Cowman et al. 2017). *P. falciparum* is the dominant malaria species in sub-Saharan Africa, where it causes the majority of malaria deaths and where Duffy negativity is highly prevalent (Hay et al. 2009; Price et al. 2020), so it is clear that *P. falciparum* does not absolutely or significantly rely on DARC to invade RBCs. However, there are many *P. falciparum* invasion ligands with receptors that have not yet been identified, including multiple orthologues of PvDBP (Cowman et al. 2017). There are also conflicting hypotheses on the origin of *P. vivax* and its relevance as a selective force for Duffy negativity (Livingstone 1984; Escalante et al. 2005; Liu et al. 2014); and some have even suggested that interaction between *P. falciparum* and *P. vivax* parasites, rather than *P. vivax* alone, may explain the near fixation of Duffy negativity in African populations (Roche et al. 2017). This leaves open the possibility that Duffy negativity may have been selected in the African population in part because it also provides some level of protection against a pathogen other than *P. vivax*.

In this study, we systematically explore the impact of Duffy blood group polymorphisms on the biophysical and membrane protein properties of human RBCs and on their susceptibility to invasion by *P. falciparum* parasites, using a combination of *in vitro* invasion assays, microscopy, and quantitative proteomics. We show, for the first time, that *P. falciparum* parasites have a clear and consistent preference for Duffy-positive RBCs. Duffy negativity was also associated with a decrease in the expression of other RBC surface proteins, including ABCA7 (ATP binding cassette subfamily A member 7). Applying an evolutionary-epidemiological model to the *in vitro* invasion data, we show that when the FYB^ES^ allele inhibits *P. falciparum* invasion at levels in line with our *in vitro* data, the frequency of the FYB^ES^ allele increases in low *P. falciparum* transmission settings, but decreases in high *P. falciparum* transmission settings. Although *P. falciparum* relies on several receptors for invasion and the exact molecular mechanism that underlies *P. falciparum’s* preference for Duffy-positive is unclear, our findings suggest that the presence of DARC receptor on the RBC could be playing either a direct or an indirect role during *P. falciparum* invasion and suggest that the expansion of *P. falciparum* may, in part, have impacted the distribution of Duffy negativity in sub-Saharan Africa.

## RESULTS

### Study design and determination of Duffy blood groups

Whole blood samples were collected in a recall-by-genotype study from 23 participants in two rounds (**Figure S1, Table S1**). In the first round (round 1), we collected samples from 20 donors in four batches (5 samples/batch), but not all four major Duffy phenotypes were represented in each batch. We therefore carried out another round of sample collection (round 2, n=16 donors) where 13 specific individuals from round 1 and 3 additional donors were recalled on the same day to generate four batches (4 samples/batch), with all four major Duffy phenotypes represented in each batch. In both rounds, the batches were processed and analysed one at a time, and the data were subsequently grouped by genotype and phenotype. The Duffy blood groups were determined both phenotypically by an immunoagglutination test and genotypically by targeted amplification and sequencing of the GATA1 and Exon 2 relevant regions of the Duffy gene (**Figure S2**), without prior knowledge of the phenotyping results. Details of these approaches and the assignment of genotypes and phenotypes to samples are described in detail in the Methods.

### *P. falciparum* parasites show a preference for Duffy-positive human RBCs

Invasion of Duffy-positive and -negative human RBCs by both *P. falciparum* and *P. knowlesi* parasites was quantified using two different *in vitro* assay approaches. In the RBC preference assay (Theron et al. 2018), synchronised schizont-stage malaria parasites were incubated together in a single well with RBCs from four different Duffy phenotypes/genotypes which had each been labelled with a different combination of fluorescent dyes (**Figure S3**). Combining all four Duffy RBC samples in the same well allows the parasites to select their “preference” from the four samples simultaneously and reduces technical variation that can occur when each blood group is present in separate wells. In the standard invasion assay, the same parasites were incubated separately with each of the four labelled Duffy RBC phenotypes in individual wells, and invasion rates compared across wells. The assays were run with both *P. falciparum* 3D7 and Dd2 strains, which have well-known differences in their utilisation of invasion receptors and are often classified as sialic acid-dependent and sialic acid-independent, respectively (Gaur et al. 2006; Spadafora et al. 2010). *P. knowlesi* parasites, which have a known reliance on the DARC receptor for invasion of human RBCs (Miller et al. 1975; Moon et al. 2013), were used as a control.

The *in vitro* assay data for round 2 samples (where each batch contained all four Duffy phenotypes) are presented in **Figure 1**, while results for round 1 samples (where each batch of samples contained only three Duffy phenotypes) are presented in **Figure S4**; results from both rounds are consistent. Both assay approaches showed that *P. falciparum* has a consistent preference for Duffy-positive RBCs (**Figure 1, Figure S4-S6**). In the preference assays, the average proportion of RBCs invaded with 3D7 was 26.7% (±5.5) in Duffy-positive (averaged across all three Duffy positive phenotypes in each assay) and 19.8% (±1.6) in Duffy-negative RBCs, and average invasion with Dd2 parasites was 27.0% (±5.6) in Duffy-positive and 19.1% (±4.1) in Duffy-negative RBCs (**Figure 1A**). In the control *P. knowlesi* parasites this preference was much more pronounced, with the average proportion of RBCs invaded being 31.3% (±7.5) in Duffy-positive RBCs and only 6.0% (±1.9) in Duffy-negative RBCs. In the invasion assay format, the average proportion of RBCs invaded with 3D7 parasites was 26.3% (±4.3) in Duffy-positive (averaged across all three Duffy positive phenotypes in each assay) and 21.1% (±2.5) in Duffy-negative RBCs, and average invasion with Dd2 parasites was 26.5% (±4.1) in Duffy-positive and 20.6% (±3.3) in Duffy-negative RBCs (**Figure 1B**). For the *P. knowlesi* control parasites, average invasion was 32.1% (±6.4) in Duffy-positive compared to only 3.8% (±0.2) in Duffy-negative RBCs. Data for each batch of round 2 samples is presented separately in **Figure S5**, and the underlying Duffy phenotypes of the grouped data in Figure 1 are in **Figure S6A-B.**

**Figure 1:**
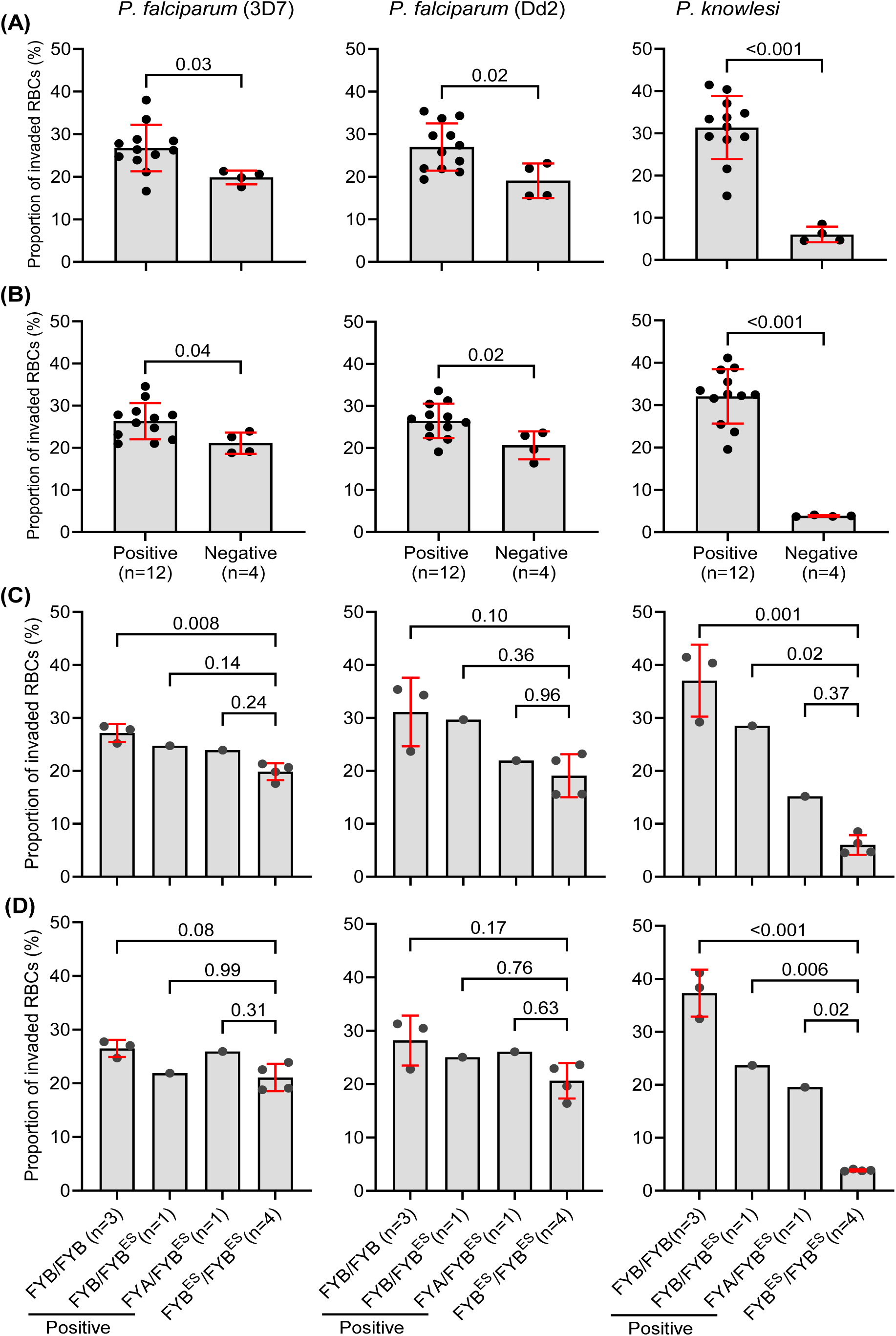
*Plasmodium falciparum* parasites preferentially invade Duffy-positive RBCs. (**A**) We used an *in vitro* preference assay where RBCs from all four major Duffy phenotypes were combined and incubated together with parasites and found a preference for the Duffy-positive RBCs by all tested parasites. (**B**) A separate assay where each of the RBCs samples was separately incubated with parasites was performed in parallel and a similar preference for the Duffy-positive RBCs was also observed. We explored the impact of heterozygosity for the FYB^ES^ (null) allele on parasite invasion by reanalysing the invasion data in A and B above; a clear gene dosage effect son *P. knowlesi* invasion was observed in both the preference (**C**) and invasion assays (**D**). For *P. falciparum* parasites, the only significant difference was between the FYB/FYB and the FYB^ES^/FYB^ES^. Statistical comparisons between panels A and B were carried out using a two-tailed t-test, while comparison for panels C and D was performed using a one-way ANOVA, followed by pairwise comparisons using Tukey’s HSD test.

The Duffy-positive RBCs used in these assays are from individuals with different underlying Duffy genotypes, which have been shown to differently affect DARC antigen expression (Woolley et al. 2000; Michon et al. 2001). To explore any link between DARC expression and *P. falciparum* invasion, we first re-analysed the invasion data to explore the impact of heterozygosity for the FYB^ES^ allele on *P. falciparum* invasion preference (the FYA^ES^ allele was not identified in any of the samples). These heterozygous samples are RBCs with either wildtype FYB or FYA in combination with the variant FYB^ES^ allele (i.e., FYA/FYB^ES^ or FYB/FYB^ES^)(**Table S2-S3**). We found that the FYB^ES^ allele had a clear dosage effect on invasion by *P. knowlesi* (**Figure 1C-D**).While there was the indication of a similar dosage effect for both *P. falciparum* strains, only the comparison between FYB/FYB and FYB^ES^/FYB^ES^ alleles reached significance in both strains and in both assay formats (**Figure 1C-D**). The limited sample number of FYB^ES^ heterozygotes limits the significance of this approach, so we therefore performed additional sets of invasion and preference assays using the same RBC samples but using combinations of RBCs designed to compare invasion efficiency between Duffy genotypes (FYB/FYB, FYA/FYB, FYA/FYA, and FYB^ES^/FYB^ES^) rather than phenotypes, as heterozygous genotypes containing the null allele express reduced levels of DARC antigens and antibody binding capacity (Woolley et al. 2000; Michon et al. 2001). Similar to the assays set up to compare invasion by phenotype, comparison of invasion by Duffy genotype showed that all parasites have a clear preference for the Duffy-positive RBCs in both assay formats (**Figure S6C-D**), with comparisons between FYB/FYB and FYB^ES^/FYB^ES^ being consistently different in all species/strains and assays, whereas differences between other genotypes were different only in some combinations (**Figure S6C-D)**. Unlike *P. knowlesi*, there is therefore no clear link between relative DARC expression and invasion by *P. falciparum*, but there is a reproducible link between absolute DARC expression (i.e., comparison of FYB/FYB and FYB^ES^/FYB^ES^) and *P. falciparum* invasion across two different assay formats and multiple biological and technical repeats.

### Biophysical properties vary between Duffy-positive and -negative RBCs

Differences in the biophysical properties of RBCs are linked to reduced parasite invasion efficiency in individuals with the rare Dantu blood group; RBCs from Dantu-positive individuals have higher surface tension and are more refractory to invasion (Kariuki et al. 2020) and inhibitory to parasite growth (Kariuki et al. 2023). To explore whether Duffy negativity causes changes in the biophysical properties of the Duffy RBCs that could explain the preferential invasion of the Duffy-positive RBCs, we used flickering spectroscopy (Yoon et al. 2009; Introini et al. 2022) to generate measurements of surface tension (resistance of the RBC membrane to stretching), bending modulus (energy required to bend the RBC membrane), viscosity (resistance to deformation), and median equatorial RBC radius of the same RBC samples used for invasion and preference assays in Figure 1. The analysis revealed minor differences in membrane tension between Duffy-positive and -negative RBCs (**Figure 2A**). Duffy-negative RBCs had a slightly higher median membrane tension (1.64*10^-6^ Nm^-1^) compared to Duffy-positive RBCs (1.39*10^-6^ Nm^-1^) (*P<0.001*); but the reverse was the case for bending modulus, with bending modulus of 1.11*10^-19^ (J) for the Duffy-negative RBCs lower relative to 1.22*10^-19^ (J) for the Duffy-positive RBCs (*P<0.001*). While this difference was significant, it was notably lower than the difference between Dantu blood group variants, where Dantu homozygotes had a median tension of 8.2*10^-7^ Nm^-1,^ and a ‘tension threshold’ above which little successful invasion occurs was proposed at 3.8 (± 2.0)*10^-7^ Nm^-1^ (Kariuki et al. 2020). Very few of the Duffy-negative RBCs fall above this threshold, suggesting that tension of Duffy-negative RBCs is unlikely to act as a significant barrier to RBC invasion. There were no statistically significant differences in viscosity and radius between Duffy-positive and Duffy-negative RBCs (*P>0.05*) (**Figure 2A**).

**Figure 2:**
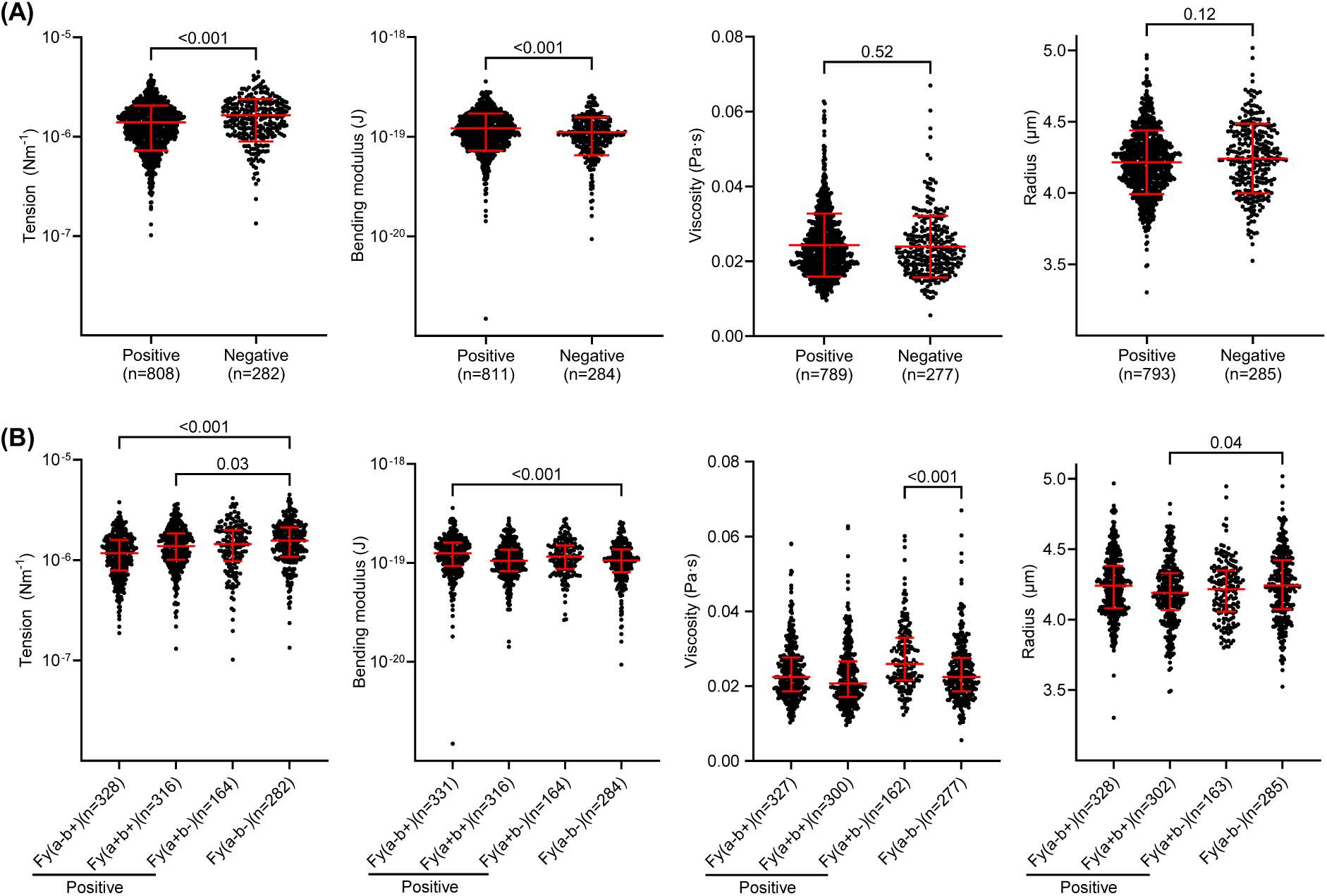
Comparing biophysical properties of Duffy RBCs as measured by flickering spectroscopy. **(A**) We measured the biophysical properties (tension, bending modulus, viscosity, and radius) of Duffy-positive (n=14) and Duffy-negative RBCs (n=5) and found that tension and bending modulus varied, but viscosity and radius were similar between the two groups. (**B)** The data in panel A was re-grouped based on Duffy RBC phenotype (Fy(a- b+)(n=6), Fy(a+b+)(n=5), Fy(a+b-)(n=3), and Fy(a-b-)(n=5)) and re-analysed, to determine the contributions of individual Duffy phenotypes, and we noted that the Fy(a-b+) and Fy(a+b+) phenotypes significantly contributed to the differences in tension and bending modulus in A above. Statistical significance for data in panel A was determined using a two-tailed Welch’s t-test, while a Kruskal-Wallis test was used to determine significance for data in panel B, followed by Dunn’s test to correct for multiple comparisons. The number of analysed RBCs, median, and interquartile range (IQR) are shown for each group.

Re-analysing the data by phenotype showed minor differences in the measured properties between the Duffy-negative and specific individual Duffy-positive phenotypes (**Figure 2B**). Notably, the differences in tension between the Fy(a-b-) and Fy(a-b+) RBCs appear to reflect the invasion preferences of the malaria parasites as determined by flow cytometry (**Figure 1**), that is, both Fy(a-b+) and Fy(a+b+) RBCs which had lower surface tension than Fy(a-b-) RBCs were more susceptible to invasion by both *P. falciparum* and *P. knowlesi* parasites. There could therefore be an association between the biophysical properties of the Duffy RBCs and parasite invasion efficiency. However, this is unlikely given the magnitude of the differences in these properties between the Duffy phenotypes. We also repeated the flickering analyses on a randomly selected subset of samples collected in the second round and noted further differences in biophysical properties between the phenotypes (**Figure S7**), suggesting that differences in biophysical properties between the Duffy phenotypes are more likely due to natural or non- genetic variations between RBC donors (Tzounakas et al. 2016), rather than caused by the Duffy GATA1 mutation.

### The relative abundance of several RBC surface proteins varied between Duffy-positive and - negative RBCs

The biophysical properties of the Duffy RBCs alone therefore do not appear to fully explain the preferential invasion of Duffy-positive RBCs by *P. falciparum* parasites (**Figure 2, Figure S7**), so we explored whether Duffy negativity could be affecting the expression of other known *P. falciparum* invasion receptors by quantifying RBC surface proteins using quantitative plasma membrane profiling (Ravenhill et al. 2019). We enriched surface proteins from 16 RBC samples representing six different Duffy genotypes and four Duffy phenotypes (**Table S1;** round 2 samples), and quantified their relative abundance in the RBC samples using mass spectrometry. In all, we identified 252 unique RBC surface proteins across all samples and compared differences in the relative abundance of the proteins between Duffy-positive and Duffy-negative phenotypes. Of the 252 proteins, 19 had significant differences in relative abundance between Duffy-positive and Duffy-negative RBCs (**Table S4, Figure 3A**). As expected, DARC was the protein most significantly depleted in Duffy-negative RBCs, thus serving as an important internal control (**Figure 3**). By contrast, we found no significant differences in the relative abundance of any known *P. falciparum* invasion receptors, including GYPA, GYPC, BSG, and Band 3 (**Figure 3B**), suggesting that the Duffy GATA1 mutation does not directly impact the expression of these receptors even in heterozygous (FYB/FYB^ES^ or FYA/FYB^ES^) individuals. However, the relative abundance of several other proteins, including ABCA7 (ATP binding cassette subfamily A member 7), was significantly reduced in Duffy-negative RBCs (**Table S4**). ABCA7, is a cholesterol transporter with known mutations that are linked to increased risk of Alzheimer’s disease, particularly in people of African ancestry (Abe-Dohmae et al. 2004; Reitz et al. 2013). ABCA7 had a remarkably consistent expression profile across all four batches of RBC samples analysed, with decreased abundance in Duffy-negative relative to Duffy-positive RBCs (*P<001*) (**Figure 3C**). Proteins enriched in Duffy-positive RBCs include MFSD2B (phospholipid transporter) (*P<0.05*), RHAG (rhesus blood group protein and ammonium transporter) (*P<0.05*), TFRC (CD71, which plays roles in iron uptake and in *P. vivax* invasion)(*P<0.05*), and several others (**Table S4**)(Tilley et al. 2010; Vu et al. 2017; Gruszczyk et al. 2018). Collectively, these data show that Duffy negativity affects the surface abundance of multiple RBC surface proteins in addition to DARC. These proteins, along with DARC, could therefore serve as potential invasion receptor candidates that explain the preference of *P. falciparum* parasites for Duffy-positive RBCs.

**Figure 3:**
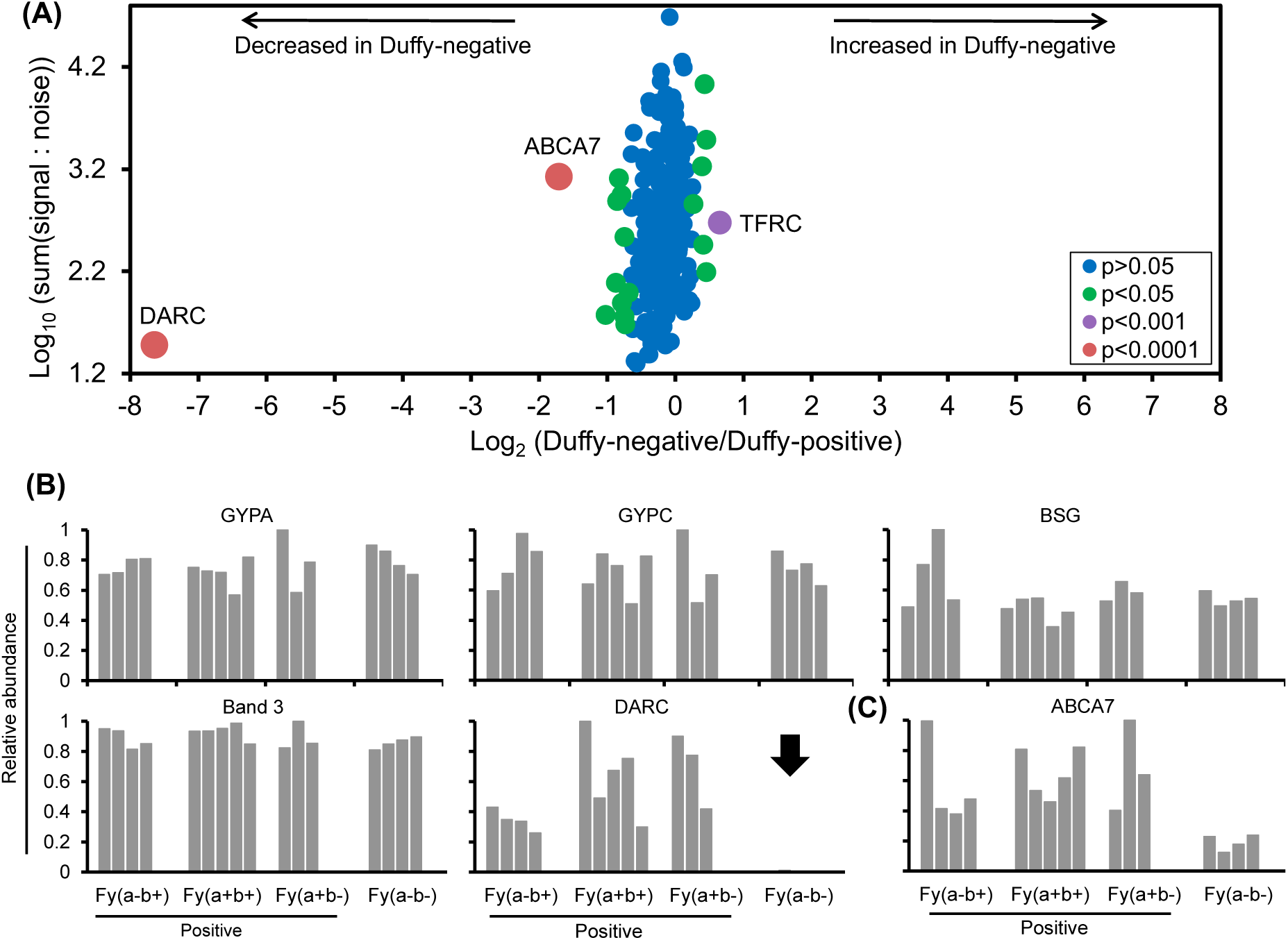
The relative abundance of surface proteins on Duffy-positive and -negative RBCs as quantified by mass spectrometry. (**A**) We used RBC plasma membrane profiling-based mass spectrometry to quantify the relative expression of surface proteins on 16 Duffy-positive and -negative RBC samples and identified 252 unique proteins. The x-axis shows the fold change, which was calculated for each protein using the signal-to-noise ratio of Duffy-negative to Duffy-positive RBCs, and the y-axis shows the summed signal-to-noise ratio of the Duffy-positive and -negative RBCs. A two-tailed t-test was used to determine the level of significance and estimate *P* values, which were corrected for multiple hypothesis testing using the Benjamini–Hochberg method. (**B**) The bar plots show the relative abundance of five RBC proteins known to be used as receptors by *P. falciparum* merozoites. (**C**) The relative abundance of ABCA7 across the Duffy blood groups. Each bar represents a sample donor and coloured according to Duffy phenotype. The black arrow shows the depletion of DARC on the RBCs of Duffy-negative individuals.

### The impact of blocking *P. falciparum* infection on the evolution of Duffy negativity

We explored the implications of our *in vitro* findings for the evolution of Duffy negativity using an evolutionary-epidemiological model. We set up a system of linked ordinary differential equations representing a genotype and age-stratified host population infected with *Plasmodium falciparum*. A similar model was previously used as a general framework to explore the evolution of malaria-blocking adaptations (Penman and Gandon 2020). Crucially, the model includes the following realistic assumptions (i) the ability of the host to develop a form of clinical immunity against the worst virulence effects of *P. falciparum*; immunity which requires a certain number of exposures to *P. falciparum* infection, and (ii) the potential for *P. falciparum* to impose not only mortality costs, but also a reproductive cost in the reproductively mature age class, a cost which is ameliorated in hosts that have already developed immunity to virulence.

If the level of *P. falciparum* erythrocyte invasion blocking observed for FYB^ES^ heterozygotes and homozygotes *in vitro* translates into the blocking of blood stage infections *in vivo*, then when *P. falciparum* transmission is relatively low (lower values of R_0_), *P. falciparum* will drive an increase in the frequency of FYB^ES^ in the population (**Figure 4**, panels A and B, blue lines). This is true for the degree of blocking observed in the invasion assays in Figure 1 for either 3D7 or Dd2, but the change in frequency tends to be greater over time for the Dd2 values (compare solid and dashed lines in Figure 4), reflecting the stronger impact on invasion preference in this line. For higher transmission levels of *P. falciparum* (higher values of R_0_), the rate of increase in FYB^ES^ is slower, and above a certain R_0_, the presence of *P. falciparum* slightly reduces the frequency of FYB^ES^ in the population (**Figure 4A** and 4B, green lines).

**Figure 4:**
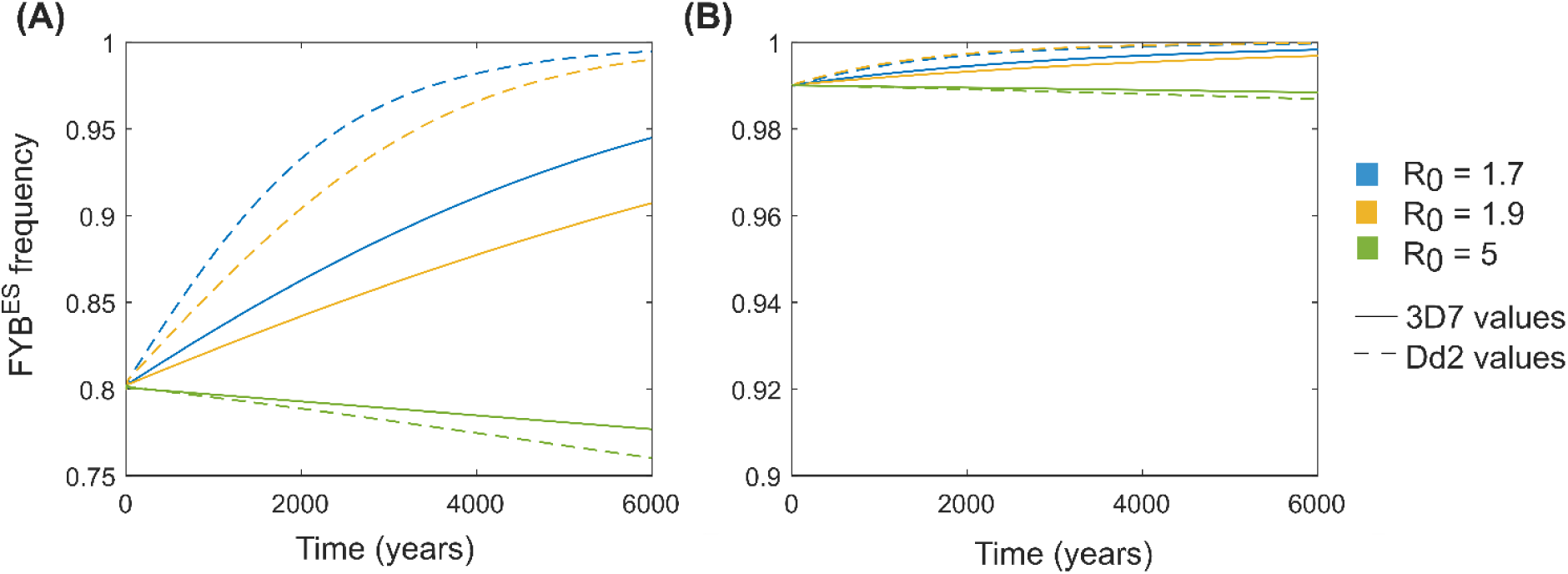
An evolutionary-epidemiological model reveals changes in FYB^ES^ frequencies induced by the presence of *P. falciparum*. Parameters are as defined in **Table 1**. (**A**) The starting frequency of FYB^ES^ is 0.8 (with genotypes in the population at Hardy-Weinberg equilibrium). (**B**) The starting frequency of FYB^ES^ is 0.99 (with genotypes in the population likewise at Hardy-Weinberg equilibrium). Different coloured lines indicate different levels of *P. falciparum* transmission (see legend and Table S6). Dashed lines indicate scenarios where we assumed that homozygosity for FYB^ES^ blocks 38.6% of infections and heterozygosity for FYB^ES^ blocks 4.5% of infections; these are the values implied by *in vitro* preference assays for Dd2 parasites (Figure 1C), assuming that the heterozygote is FYB^ES^/FYB. Solid lines indicate scenarios where we assumed that homozygosity for FYB^ES^ blocks 26.9% of infections and heterozygosity for FYB^ES^ blocks 8.86% of infections; these are the values implied by *in vitro* preference assays for 3D7 parasites (Figure 1C).

This relationship between *P. falciparum* transmission and FYB^ES^ behaviour emerges because, in keeping with the known properties of *P. falciparum*, we have assumed that protective immunity to the most significant clinical effects of *P. falciparum* can be gained after relatively few infections. FYB^ES^ containing genotypes therefore have the greatest *relative* advantage in settings where few people are likely to gain full immunity to malaria virulence before reproductive maturity (i.e., low transmission settings). This phenomenon is fully explored in (Penman and Gandon 2020). Note that *P. vivax* (the parasite which is widely believed to have first elevated FYB^ES^ frequencies) does not display the same severe syndromes associated with immune naivity as does *P. falciparum*, thus we do not expect selection from the two parasites to operate in the same way.

The impact of *P. falciparum* on FYB^ES^ frequencies over the likely timescales when *P. falciparum* has acted as a selective pressure on human populations (within the last ∼10 thousand years and potentially linked to the advent of agriculture ∼6000 years ago (Rich et al. 1998; Sundararaman et al. 2016)) depends on the starting frequency of FYB^ES^. If FYB^ES^ is at a frequency of 0.8 when selection from *P. falciparum* begins (**Figure 4A**), much greater changes in allele frequencies occur over 6000 years than if FYB^ES^ is at a frequency of 0.99 when *P. falciparum* arrives (**Figure 4B**). These starting frequencies of FYB^ES^ were chosen to be representative of FYB^ES^ frequencies currently seen in the Central African Republic (∼0.8) and West Africa (>0.99) (**Table S6**).

### The impact of *P. falciparum* on Duffy negativity may be most apparent in central Africa

Given the negative relationship we predict between *P. falciparum* transmission intensity and FYB^ES^ frequency (**Figure 4**), it would be useful to have estimates of historical *P. falciparum* transmission intensities and investigate their relationship with FYB^ES^. The frequency of the sickle cell allele (HbS), which offers profound protection against severe *P. falciparum* malaria, is a potential proxy for the past intensity of *P. falciparum* transmission in African populations. We obtained FYB^ES^ and HbS frequencies from different African populations published by MalariaGEN (Malaria Genomic Epidemiology Network et al. 2019), and supplemented these with anthropological studies that measured both Duffy negativity and sickle cell frequencies (**Table S6**). These anthropological studies were selected from the literature database that supports Howes *et al*’s global map of Duffy negativity (Howes et al. 2011). We assigned populations to four geographical subgroups, based on their locations relative to the likely origin of the Bantu expansion (**Table S6** and **Figure 5A**). This was because the Bantu expansion potentially had a substantial impact on FYB^ES^ frequencies in different regions. The northern group of populations and the southeastern group all had extremely high frequencies of FYB^ES^ (>0.99, **Table S6, Figure 5B**). The southcentral group had the lowest frequencies (between 0.8 and 0.9), and the southwestern group had FYB^ES^ frequencies between 0.8 and 1. It seems likely that this pattern is a consequence of different patterns of population admixture in different regions during and after the Bantu expansion (Fortes-Lima et al. 2024).

**Figure 5:**
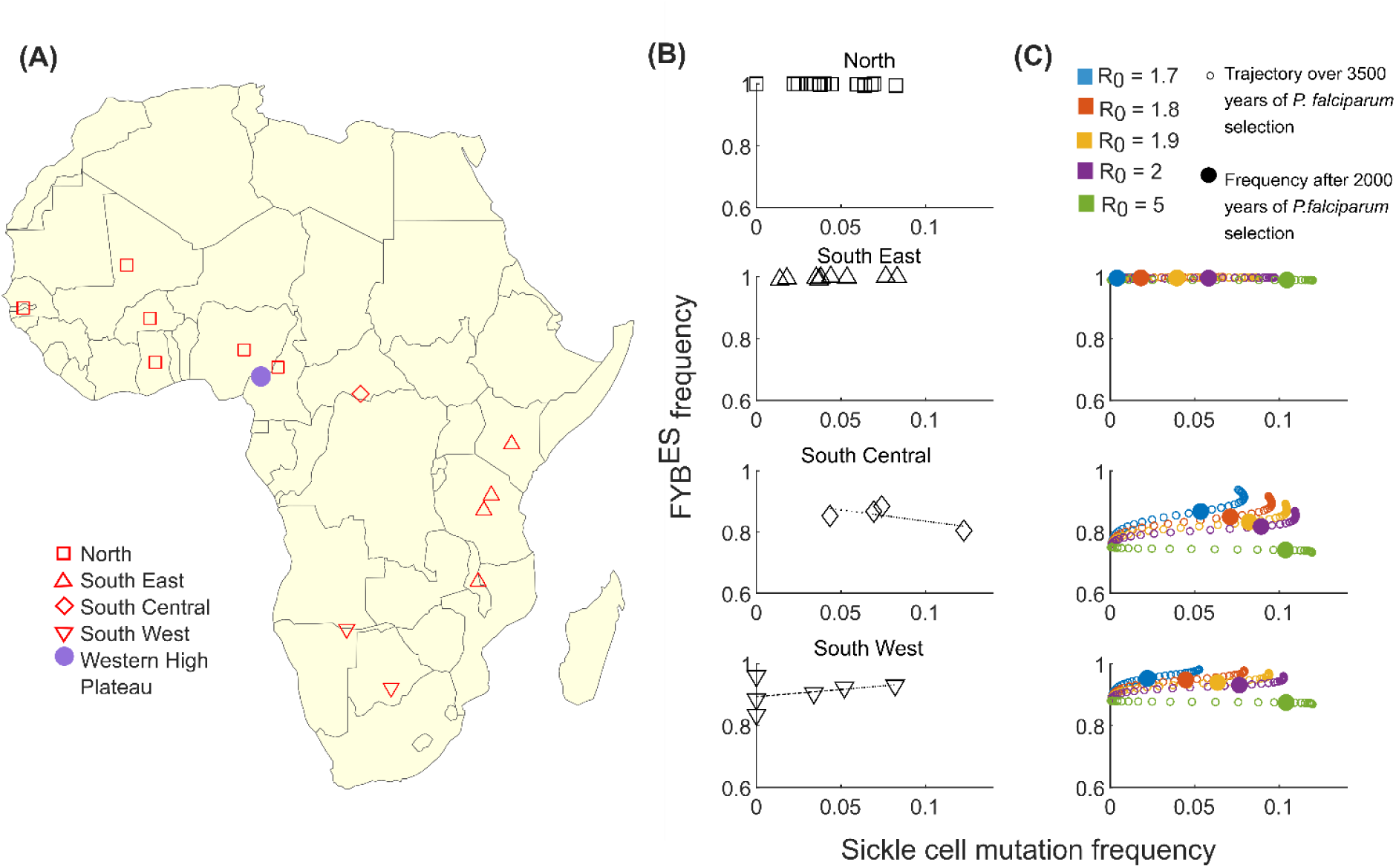
Duffy negativity and likely historical selection from *P. falciparum* in African populations. (**A**) The locations of the populations studied (Table S6). For populations where the specific location of the study is available, this has been used, otherwise we used the centroid of the country where the relevant population was sampled. The shape of each marker indicates how we assigned geographical subgroups. The Western High Plateau of Cameroon is indicated on the map as a potential origin point for the Bantu expansion. (**B**) The relationship between the frequency of the sickle cell mutation (a proxy for the strength of historical *P. falciparum* selection) and the frequency of the FYB^ES^ mutation in the African populations listed in Table S6, separated by geographical subgroups. (**C**) The results of our evolutionary-epidemiological model, simulating both the FYB^ES^ mutation and the sickle cell mutation. We ran simulations for 3500 years, in keeping with the potential timing of the Bantu expansion into South Central and South Western Africa. We tested 5 different levels of malaria transmission (different colours). Small unfilled circles indicate combinations of sickle cell and FYB^ES^ frequencies observed from time point zero up until time point 3500 years. The frequencies obtained after 2000 years of evolution are also highlighted by larger filled circles. The starting frequency of the sickle cell mutation was always 0.0005. We tested three different starting frequencies for FYB^ES^ in Figure 5C: 0.99 (top panel), 0.75 (middle panel), and 0.88 (bottom panel). These values were chosen for illustrative purposes, to show that the model can capture combinations of FYB^ES^ and sickle cell frequencies that are similar to those observed in different regions of Africa.

If sickle cell is a proxy for past *P. falciparum* selection (such that higher sickle cell frequencies indicate more intense historical exposure to *P. falciparum*), and FYB^ES^ impedes *P. falciparum* invasion in the way we observe, our model predicts a negative relationship between sickle cell frequencies (i.e. *P. falciparum* intensity) and FYB^ES^ frequencies. As shown in Figure 5B, we only find such a relationship for the southcentral region. However, when we expand our model to include the sickle cell mutation alongside the FYB^ES^ mutation, we can see why this should be the case. The population genetics of the DARC locus implies that FYB^ES^ increased in frequency, with a very strong selection coefficient, at some point during the last 50 thousand years (McManus et al. 2017). This means that it is possible that FYB^ES^ reached frequencies close to fixation in some populations in sub-Saharan Africa before *P. falciparum* became a selective pressure ∼6000-10000 years ago (Rich et al. 1998; Sundararaman et al. 2016). When we simulate both FYB^ES^ and sickle cell under selection from *P. falciparum*, starting with FYB^ES^ at a very high frequency but sickle cell at a low frequency, varying levels of *P. falciparum* intensity easily generate a realistic range of sickle cell frequencies within a plausibly short time frame (**Figure 5C**, top panel), but the frequency of FYB^ES^ barely moves away from its very high frequency, no matter what the intensity of *P. falciparum.* This creates a pattern that is consistent with the lack of relationship between sickle cell and FYB^ES^ in the North and South Eastern regions (**Figure 5B**).

For the South Central region, by contrast, let us assume that (due to different levels of admixture between FYB^ES^ high and low frequency populations), FYB^ES^ was at a lower frequency when selection from *P. falciparum* began. A range of levels of *P. falciparum* selection intensity can still generate a range of different sickle cell frequencies (**Figure 5C**, middle panel), with the highest levels of sickle cell observed at the highest *P. falciparum* transmission levels. However, as explored in Figure 4, FYB^ES^ is most advantaged at *lower P. falciparum* transmission intensities, and its frequency may be suppressed when *P. falciparum* transmission is high. Their contrasting relationships with *P. falciparum* transmission intensity mean that a negative relationship emerges between FYB^ES^ and sickle cell frequencies in our model, which matches the pattern seen in the South Central data (**Figure 5B**).

South Western populations exhibit a weak positive correlation between FYB^ES^ frequencies and sickle cell (**Figure 5B**, bottom panel), which contradicts our prediction. However, when we set the starting frequency of FYB^ES^ to be 0.88, the pattern of FYB^ES^ and sickle cell mutation frequencies obtained for R_0_ values between 1.7 and 2 is within a similar range to the actual frequencies observed in Southwestern populations, and this particular set of starting conditions is only associated with a relatively weak relationship between FYB^ES^ and sickle cell frequencies in our model (**Figure 5C**, bottom panel). The lack of data for populations in Central and South Western Africa limits our ability to draw strong conclusions. Future studies should focus on these regions in order to determine if there truly is any negative relationship between past *P. falciparum* selection (measured by sickle cell frequency or another proxy) and FYB^ES^ frequency, as predicted by the results we present here.

## DISCUSSION

The high frequency of the FYB^ES^ allele in West and Central Africa has historically been attributed to selection by *P. vivax,* which has a significantly reduced ability to invade Duffy-negative RBCs and is, as a result, less prevalent in sub-Saharan Africa (Miller et al. 1976; Howes et al. 2011). However, there is increasing evidence suggesting that individuals who are Duffy-negative are not completely protected against *P. vivax* (Ménard et al. 2010; Twohig et al. 2019). This, combined with the geographic distribution and prevalence of *P. falciparum* in sub-Saharan Africa (Weiss et al. 2019), raises the possibility that the FYB^ES^ allele may have been positively selected because it protects not only against *P. vivax* but also against other frequently found pathogens in the region, possibly including *P. falciparum*. In this study, we used a recall-by-genotype design and a combination of *in vitro* assays, quantitative proteomics, and flickering spectroscopy approaches (Yoon et al. 2009; Theron et al. 2018; Ravenhill et al. 2019), to investigate the impact of Duffy blood group polymorphisms on *P. falciparum* invasion and the RBC itself. We also used an evolutionary-epidemiological model to explore the effect of Duffy negativity blocking *P. falciparum* invasion on the frequency of the FYB^ES^ allele in different *P. falciparum* transmission settings.

We found that *P. falciparum* parasites (both 3D7 and Dd2 strains, which use different invasion pathways (Gaur et al. 2006; Spadafora et al. 2010)) have a preference for Duffy-positive over Duffy-negative RBCs. The preference for Duffy-positive RBCs was consistent regardless of whether each of the *P. falciparum* strains was co-incubated together with all four Duffy RBC groups in the same well in a preference assay (**Figure 1A, Figure S4**) or each Duffy RBC group was incubated separately with each of the *P. falciparum* strains in an invasion assay (**Figure 1B**). We also assessed invasion using an independent methodology, video microscopy (**Figure S8**), which produced results that were consistent with a preference of *P. falciparum* for Duffy-positive RBCs, although the low throughput nature of these assays means that the video data did not meet statistical cut-offs for significance. It is of course abundantly clear that DARC is not absolutely required for *P. falciparum* invasion, given the widespread prevalence of *P. falciparum* in sub-Saharan Africa, where Duffy-negativity is near-ubiquitous (Weiss et al. 2019). However, our data show that DARC does have an impact on *P. falciparum* invasion, and adds a significant new element to our understanding of the host factors involved in *P. falciparum*-RBC interactions.

One of the limitations of this study is that protection against *P. falciparum* is multifactorial, and it is not possible to screen for all potentially protective polymorphisms. Samples were screened for HbS, and none of the samples carried this allele. We also typed the samples for ABO blood groups, and regression analysis controlling for ABO blood group had no significant impact on the invasion data – **Table S5**). While it is formally true that other untested polymorphisms could be confounding, the effect was consistent across all donors (**Figure S5**), and it seems unlikely that all donors would carry such mutations. The sample size (n=23) is also a limitation of the study. However, the complex experimental design (matched samples from multiple Duffy blood groups compared at the same time in technical triplicates and biological duplicates) limits scaling to much larger numbers of samples, but also increases the reliability and robustness of the comparisons and findings.

There are two logical mechanistic explanations for our finding - either a *P. falciparum* invasion ligand directly interacts with DARC as a receptor during invasion; or Duffy negativity may indirectly impact invasion by modulating biophysical properties of the RBC, or by impacting the expression of one or more other RBC receptors. A direct effect of Duffy negativity on invasion is certainly possible. Unlike *P. vivax* and *P. knowlesi* which both rely almost exclusively on the DARC receptor for invasion (Miller et al. 1975; Miller et al. 1976), *P. falciparum* uses multiple receptors for invasion (Salinas and Tolia 2016). Both *P. vivax and P. knowlesi* bind to the DARC receptor through the Duffy-Binding Protein (DBP) (Batchelor et al. 2014; Gunalan et al. 2016). The *P. falciparum* genome contains multiple homologues of Pv/PkDBP, collectively referred to as the erythrocyte-binding antigen (*Pf*EBA) family, members of which play key roles in *P. falciparum* invasion and bind to glycophorin receptors on the RBC surface (Gilberger et al. 2003; Maier et al. 2003). There are still however members of the EBA family, as well as members of a different invasion ligand family, the *P. falciparum* reticulocyte binding-like (*Pf*Rh) family, whose RBC receptors are unknown (Tham et al. 2012; Salinas and Tolia 2016), raising the possibility that DARC could be a receptor for one of these ligands. Supporting this direct interaction hypothesis, we found that the FYB^ES^ allele had a clear gene dosage effect on *P. knowlesi* invasion, a species that relies extensively on DARC for invasion (**Figure 1C-D**), and an indication of a similar effect on *P. falciparum* invasion, as only the comparison between FYB/FYB and FYB^ES^/FYB^ES^ alleles reached significance in both strains and in both assay formats (**Figure 1C-D, Figure S6C-D**). This observation is consistent with an increased expression of DARC in FYB/FYB RBCs (phenotype: Fy(a-b+)) relative to other Duffy blood groups (Woolley et al. 2000; Michon et al. 2001; Ménard et al. 2010; King et al. 2011). In line with these findings, we observed that the Fy(a-b+) RBCs were associated with higher *P. knowlesi* invasion compared to the Fy(a+b-) RBCs (**Figure 1, Figure S4B**). Our findings therefore add weight to the possibility that DARC is involved directly as a receptor during *P. falciparum* invasion, and that the relative abundance of DARC on the RBC surface and its blood group antigenic variations are linked to invasion efficiency.

An indirect effect of Duffy gene mutations on the invasion of Duffy-negative RBCs by *P. falciparum* is also possible. The human RBC plasma membrane is simpler than the plasma membrane of most eukaryotic cells, but it still contains hundreds of proteins, some of which are known to form complexes (Bruce et al. 2003; Kodippili et al. 2020). For example, the DARC glycoprotein is part of a multi-protein complex (erythrocyte P55, GYPC, XK, Kell, Rh, Duffy, and Band 3) which is linked to the RBC cytoskeleton via Protein 4.1R (Salomao et al. 2008), raising the possibility that the absence of DARC could have an indirect effect on proteins within and outside the complex. We therefore used RBC surface expression profiling to systematically define the RBC surface of Duffy-negative RBCs for the first time. We found no significant differences in the relative abundance of proteins within the DARC-associated complex (**Figure 3, Table S4**), nor was the relative abundance of any established *P. falciparum* invasion receptors different between Duffy-positive and -negative RBCs (**Figure 3**). However, the abundance of several other proteins which have not previously been linked to malaria invasion or resistance or susceptibility varied significantly (**Table S4**). ABCA7 in particular showed a remarkably consistent decrease in abundance in Duffy-negative RBCs (**Figure 3**). Mutations in a related transport protein, ABCB6, which encodes the Langereis (Lan) blood group has previously been linked to *P. falciparum* resistance (Egan et al. 2018). It is therefore possible that ABCA7 (or indeed any of the other proteins that are reduced in abundance (**Table S4**)) is acting as a receptor for *P. falciparum*, and that the impact of Duffy negativity is indirect only, by reducing the expression of this receptor.

Alternatively, Duffy negativity could in theory be operating indirectly by changing the biomechanical properties of RBCs, rather than modulating the relative expression of specific receptor(s). Duffy-negative RBCs had higher median membrane tension and lower bending modulus than Duffy-positive RBCs (**Figure 2**). However, while higher tension prevents invasion as seen in the Dantu RBCs (Kariuki et al. 2020), the increase in RBC tension in Duffy-negative RBCs is lower than that seen in beta thalassaemic and Dantu homozygous RBCs, respectively (Kariuki et al. 2020; Introini et al. 2022). The minor increase in tension is also counteracted by a decrease in bending modulus, as previously shown in beta thalassaemic RBCs (Introini et al. 2022), suggesting that tension is not fully responsible for the differences in invasion preference. Furthermore, membrane tension and radius have a linear inverse relationship (i.e., tension increases as radius decreases)(Popescu et al. 2006), thus the absence of differences in radius between the Duffy-positive and -negative RBCs (**Figure 2A**), also implies that membrane tension alone does not explain the differential invasion of the Duffy RBC groups. Individual donor data suggest that the differences in biophysical properties between the Duffy RBC groups are likely due to natural or non-genetic variations between donor RBCs (Tzounakas et al. 2016; Roubinian et al. 2022).

Given the majority of people in West and Central Africa are Duffy-negative (Howes et al. 2011), and most cases of *P. falciparum* malaria in sub-Saharan Africa are by definition uncomplicated (Bassat et al. 2008; Geleta and Ketema 2016), a minor protective advantage conferred by Duffy negativity against *P. falciparum*, as suggested by our invasion data, may have been previously overlooked. The overwhelming selective force that *P. falciparum* has exerted on the human genome, and the high levels of *P. falciparum* malaria mortality in the pre-antimalarial era, makes it important to consider whether even a minor impact of Duffy negativity on *P. falciparum* infection could have impacted the distribution of the FYB^ES^ allele. Our evolutionary-epidemiological model shows that partially blocking *P. falciparum* infection is most advantageous at relatively low transmission levels of *P. falciparum* itself. We therefore propose that the impact of *P. falciparum* on FYB^ES^ frequency would be greatest in low transmission intensity settings. We further show that *P. falciparum* can drive substantial changes in FYB^ES^ frequency over a realistic 6000 year period, but only if FYB^ES^ was not already at a very high frequency (**Figure 4**). This raises the intriguing possibility that in regions where malaria transmission in general was less intense (and hence perhaps FYB^ES^ was not elevated to such a high level by *P. vivax*), the arrival of relatively low intensity *P. falciparum* transmission could have been the final factor that pushed FYB^ES^ up to fixation.

We must acknowledge that our model is a highly abstracted simulation of malaria epidemiology, and is limited by the fact that most of its parameters (**Table S7**) are plausible estimates, as opposed to rates that can easily be measured. The best evidenced parameters we have used are the protective effect of sickle cell heterozygosity against clinical and severe malaria, which have been established across multiple studies (Taylor et al. 2012). The fact that our model can reproduce plausible sickle cell frequencies within plausible timescales for human evolution is, therefore, the best evidence of our model’s biological relevance. In terms of modelling the impact of FYB^ES^ genotypes, we have had to make the strong assumption that the infection blocking effect observed *in vitro* translates directly to the blocking of infections *in vivo*. The blocking of x% of invasions *in vitro* may well not translate exactly to the blocking of x% of infections *in vivo* , but the broad principle that *in vitro* inhibition of invasion translates into *in vivo* blocking of infections is borne out by a malaria challenge study in which individuals with the Dantu blood group (highly *P. falciparum* invasion blocking *in vitro*) were also protected from infection in the challenge study *in vivo* (Kariuki et al. 2023).

The spread of the FYB^ES^ mutation in sub-Saharan Africa is a textbook example of recent positive selection acting on the human genome, but the exact nature of the selective pressure has yet to be elucidated. Selection from *P. vivax* remains the leading explanation, but, as we demonstrate for the first time here, Duffy negativity can also impact infection with *P. falciparum*. A full understanding of the history of FYB^ES^ therefore requires consideration of both parasite species.

## Acknowledgements

This work was supported by the Wellcome Trust (JR: Wellcome Investigator Award 220266/Z/20/Z) and a Wellcome Senior Clinical Research Fellowship to MPW (108070/Z/15/Z). We are grateful to the Gates Cambridge Trust for supporting BM’s PhD research, and the following groups and individuals for their support and expertise: Weekes Lab at CIMR (Leah Hunter; plasma membrane profiling); CIMR Proteomics facility (John Suberu); CIMR flow cytometry facility (Reiner Schulte and Gabriela Grondys-Kotarba; flow cytometry); Cicuta group (flickering spectroscopy); Gavin Band at the University of Oxford (MalariaGEN data); and Rayner lab (Theresa Feltwell).

## Author Contributions

Conceptualisation: JCR; Formal analysis: BM, VI, RA, and BSP; Investigation: BM, VI, and BSP; Resources: JCR, JR, MR, PC, and RA; Writing-original draft: BM, VI, RA, MPW, BSP, and JCR; Writing-review & editing: BM, VI, JR, MR, RA, PC, MPW, BSP, and JCR; Funding acquisition: JCR; Supervision: JCR, MPW. All authors reviewed and approved the final manuscript.

## Data Availability

The mass spectrometry proteomics data have been deposited to the ProteomeXchange Consortium via the PRIDE partner repository, under the dataset identifier PXD065766 and project name, ‘Invasion preferences suggest a possible role for *Plasmodium falciparum* parasites in the expansion of Duffy negativity in West and Central Africa’. All data supporting the findings of this study are available within the manuscript and supplementary information.

## METHODS

### Study design and participants

Whole blood samples were collected from 23 participants between the ages of 20 to 59 in two rounds (**Figure S1**). In round 1, 8ml of whole blood was collected from 20 participants; and in round 2, another 8ml of blood was collected from 16 participants (13 of the 20 participants from round 1 were re-called, and three new participants were added). In both rounds, samples were received in batches of five (round 1) or four (round 2). In round 1, only three of the four major Duffy blood groups were represented in each batch, whereas all four Duffy blood groups were represented in each of the batches of round 2, this was useful for the erythrocyte preference assays as it meant all blood groups could be mixed and incubated together with malaria parasites in the same well. The participants were not previously genotyped for Duffy blood group antigens, thus we used ethnicity to estimate Duffy phenotype prior to sampling, as the frequency of Duffy blood group phenotypes varies both ethnically and geographically (Howes et al. 2011). All 23 study participants are part of a database of research blood donors that is maintained by Research Donors Ltd, UK, who also carried out donor recruitment and acquisition of informed consent for study participation (Ethical approval-Fulham NHS REC Reference: 20/LO/0325).

### RBC sample processing and storage

The blood samples were collected in a vacutainer with acid-citrate-dextrose (ACD) anticoagulant as a preservative. RBCs were separated from whole blood by centrifugation and washed in Roswell Park Memorial Institute (RPMI) 1640 medium (0.4g/L of Albumax II, and 2.0 g/L of Sodium bicarbonate without L-Glutamine and gentamicin (incomplete medium)). A Giemsa-stained thin blood smear was prepared to check the purity of the isolated RBCs by microscopy, and the purified RBCs were diluted to 50% hematocrit (HCT) and kept at 4⁰C for use in parasite invasion assay within 24 hrs after collection or preserved in Glycerolyte 57 solution and stored at -80°C for future use.

### Malaria parasite cultures

Both *P. falciparum* and *P. knowlesi* parasites were routinely cultured in human O+ RBCs (however, other ABO blood groups were used when O+ blood was not available) obtained from NHS Blood and Transplant (NHSBT, Cambridge, UK; ethical approval-University of Cambridge REC HBREC.2019.40 and NHS REC 20/EE/0100). Before culture, the RBCs were washed (4ml of O+ RBCs were suspended in 7ml of RPMI 1640, centrifuged twice at 3000 rpm for 5 mins at room temperature (RT)). The two *P. falciparum* strains were cultured at 4% HCT in RPMI 1640 medium containing 0.4g/L of Albumax II, 2.0 g/L of sodium bicarbonate, and supplemented with 0.292g/L of L-Glutamine and 0.025g/L gentamicin (complete medium); while the *P. knowlesi* parasites were cultured at 2% HCT using a separate complete RPMI 1640 medium formulated as above but supplemented with 10% heat-inactivated Horse Serum (Thermo Fisher). All parasites were cultured in canted-neck cell culture flasks with vented caps (Corning®) and maintained at 37°C under 1% O2, 3% CO2, and 96% N2. The culture media was regularly changed and fresh RBCs added to maintain healthy parasite growth. Parasitaemia and parasite growth were regularly checked by microscopic examination of a thin blood smear prepared from a 3-4μl aliquot of the cultures.

### Parasite synchronisation

The *P. falciparum* parasites were synchronised at ring stages using 5% D-Sorbitol, then incubated for an additional 24 hrs to ensure that parasites were at the schizont stages (confirmed by Giemsa staining and light microscopy) prior to setting up the invasion assays. Briefly, cultures were transferred into a 15 ml Falcon tube and centrifuged for 5 mins at 1100 rpm. The pellet containing both parasitised and non-parasitised RBCs was re-suspended in 10x the pellet volume of 5% D-Sorbitol, vortexed for 15 seconds, and incubated at 37⁰C in a water bath for 5 mins. The pellet was then washed in 12 ml of incomplete RPMI 1640, before an appropriate volume of complete media and fresh RBCs were added to maintain the culture at 4% HCT. The synchronised ring-stage parasites were incubated for 24 hrs to reach the schizont stage for use in the invasion assays. In contrast to *P. falciparum*, the *P. knowlesi* parasites were synchronised by density gradient separation using Nycodenz solution (10 mM HEPES and 27.6g of Histodenz, filter-sterilised, pH 7). First, the *P. knowlesi* cultures were pelleted by centrifugation (1000 rpm for 5 mins), before the pellet was re-suspended in residual supernatant and gently layered over a pre-warmed 55% Nycodenz solution, and centrifuged for 11 mins at 1500 rpm (acceleration 3, brake 1). The layer of late schizonts was washed in pre-warmed RPMI 1640 and used in the invasion assays, while the layer of ring-stage parasites and uninfected RBCs was immediately resuspended in appropriate volumes of culture media and RBCs and re-cultured.

### Duffy blood group phenotyping and genotyping

The blood samples were phenotyped by immunoagglutination using antibodies for the Duffy blood group antigens (Anti-Fy^a^ (Monoclonal), (Cat No: BG-FYA2), and Anti- Fy^b^ (Polyclonal), (Cat No: BG-FYBH2), Rapid Labs Ltd, Colchester, CO7 8SD), and genotypes were independently confirmed by sequencing the GATA1 and Exon 2 regions of the Duffy (FY) gene after amplification by polymerase chain reaction (PCR). Genomic DNA was extracted from the samples using a Qiagen DNeasy Blood & Tissue Kit (Cat No: 69504) following the kit manufacturer’s user guide with minor modifications to increase yield and concentration. Previously described hemi-nested PCR primers were used to amplify the GATA1 and Exon 2 regions (Ménard et al. 2010); and amplicons were detected by gel electrophoresis (5μl of SYBR Safe EtBr in 1% Agarose in TAE buffer, 5μl of PCR product, 100volts for 30mins), prior to being purified and sequenced. We used Chromas software (version 2.6.6) to view the trace files and Clustal Omega to align the sequences to the reference gene (ENSG00000213088). Established references were used for assigning the Duffy genotypes and phenotypes (Wurtz et al. 2011; ISBT, 2021). In round 1, all phenotyping and genotyping results were in concordance, except for one sample. For the discordant sample (**Table S2**, sample 413), the phenotyping showed that the donor has the Fy(a+b-) phenotype; however, a closer examination of the DNA sequence revealed that the participant has the rare FYA/FYX2 genotype with a predicted phenotype of Fy(a+b^weak^). This rare phenotype is due to the presence of three mutations at positions 125G>A/G, 265C>T/C, and 298G>A/G in Exon 2, which result in weak expression of the Fy^b^ antigen on the RBC surface; these mutations make the Fy^b^ antigen undetectable by the anti-Fy^b^ antibodies used in standard agglutination tests (Olsson et al. 1998). The discordant sample was assigned the phenotype predicted by the sequencing results, and the RBCs were not used in any of the assays described in this study. In round 2, all phenotyping and genotyping were also in concordance except for one sample (**Table S2**, sample 559; one of the three additional samples collected in round 2). The single discordant sample in round 2 was also immunophenotyped as Fy(a+b-); however, analysis of sequences from the GATA1 and Exon 2 regions of the gene showed that the donor has no GATA1 mutation (i.e., -67 T>T; Duffy-positive) and is heterozygous for the co-dominant FYA and FYB alleles (125G>G/A) (**Figure S2B**). The presence of a 298G>G/A substitution also confirmed the existence of both alleles at the locus. We re-extracted genomic DNA from the entire batch that contained the discordant sample and repeated both the PCR and Sanger sequencing, but the results did not change. The discordant sample was assigned the genotype FYA/FYB (phenotype: Fy(a+b+)) for all subsequent analyses. All samples were also genotyped for the malaria-protective sickle cell trait (HbAS) by PCR-RFLP as previously described (Caroca and de Lima 2016), to negate the potential confounding effect of the trait on donor RBCs, this is particularly important for the *in vitro* parasite invasion assays. No samples carried the HbS alelle.

### *In vitro* erythrocyte preference and invasion assays

We used two flow cytometry-based assays in parallel to quantify malaria parasite invasion preference for the Duffy blood groups. In the first assay, schizont-stage parasites were incubated together with RBCs from all four Duffy blood groups, each of which had been separately labelled with either 1µM or 10µM of CellTrace^TM^ Far Red (CTFR), or 10µM of Oregon Green 488 (OG), or a combination of both dyes (1µM of CTFR and 10 µM of OG) (Theron et al. 2018). The fluorescent dyes were rotated between Duffy blood groups, so no one blood group is always labelled with the same dye. The labelled Duffy-positive and -negative RBCs were all co-incubated together in the same wells with *P. falciparum* parasites at 1-2% starting parasitaemia in the ratio 1:1:1:1:1 (i.e., 20µL each of Fy(a-b+), Fy(a+b+), Fy(a+b-), Fy(a-b-), and synchronised culture; 100µl/ well) (**Figure S3A**). In the second assay format, we co-incubated the same parasites with labelled RBCs of only one Duffy blood group at a time in the ratio 1:1 (50µl of labelled RBCs: 50µl of synchronised culture;100µl/well). In both assays, each RBC samples was tested in triplicate with both 3D7 and Dd2 *P. falciparum* strains, and *P. knowlesi* parasites (control), and each assay was repeated as biological duplicates on a different day with the same RBC samples and parasites. The two *P. falciparum* strains (3D7 and Dd2) were tested in the assays because they are widely used lab strains that originate from Africa and Southeast Asia, respectively (van Schalkwyk et al. 2013; Preston et al. 2014). In both assay formats, invasion was allowed to proceed for 24 hrs at 37⁰C, after which parasitised RBCs were labelled with 2µM of Hoechst 33342 - a fluorescent dye which binds to DNA, and the percentage of parasitised RBCs in each labelled Duffy RBC group was determined by flow cytometry.

The samples were examined with a 488nm blue laser, a 640nm red laser, and a 405nm violet laser on an Attune™ NxT Acoustic Focusing Cytometer. OG was excited by the blue laser and detected by a 450/50 filter, CTFR was excited by the red laser and detected by a 670/14 filter, and Hoechst 33342 DNA dye was excited by the violet laser and detected by a 530/30 filter. For each sample, 50,000 events were recorded and the flow cytometry standard (FCS) files were analysed using FlowJo (version v10.7.2). Singlets were gated using forward scatter (FSC-H vs FSC-A), and the populations of interest (i.e., parasitised and non-parasitised RBCs for each Duffy blood group) were gated using the fluorescent intensities of the respective dyes (**Figure S3B**). The proportion of parasitised RBCs in each Duffy RBC group was determined by dividing the number of parasitised RBCs in each blood group by the total number of RBCs in the group. This proportion was then normalised by dividing the percentage (% ) parasitaemia for each group by the total % parasitaemia of all four Duffy blood groups, to allow for comparison of invasion across the different Duffy RBC groups tested.

### Video microscopy

We used video microscopy to quantify key invasion parameters during merozoite-RBC interaction in real-time, as previously described (Kariuki et al. 2020). Briefly, parasitised RBCs (late-stage schizonts) were isolated by magnetic separation using LD columns (Miltenyi Biotec) and re-suspended in complete medium with each Duffy RBC group at 0.2% HCT right before analysis. The RBCs were loaded into a separate SecureSeal hybridisation chamber (Sigma), one RBC sample at a time, and analysed under the same experimental conditions. A custom-built temperature-control system was used to maintain an optimal culture temperature of 37°C during the experiment. The samples were placed in contact with a transparent glass heater driven by a PID temperature controller in a feedback loop, with the thermocouple attached to the glass slide. A Nikon Eclipse Ti-E inverted microscope was used with a Nikon 60x Plan Apo VC numerical aperture (NA) 1.40 oil-immersion objective, kept at physiological temperature with a heated collar. The motorised functions of the microscope were controlled by custom software, and focus was maintained throughout the experiments using the Nikon Perfect Focus system. Each sample was recorded for 2 hrs, and images were acquired in bright- field with a red filter using a CMOS camera (model GS3-U3-23S6M-C, Point Grey Research/FLIR Integrated Imaging Solutions (Machine Vision), Ri Inc.) at 4 frames per second (fps), with a pixel size of 0.0986μm.

Each sample was recorded for 2 hrs, and one video was recorded for each egress–invasion event, from a few minutes before schizont rupture until the end of echinocytosis, at around 20 mins after egress. For each video, we quantified the number of merozoites that contacted the RBCs, and either invaded successfully or failed to invade. We defined the ‘parasite invasion efficiency’ or proportion of invasion, as the fraction of merozoites that contacted and successfully invaded RBCs divided by the number of all merozoites that contacted nearby RBCs post-egress, expressed as a percentage; the definition takes into account the fact that multiple invasions of the same RBC can occur. We also determined the degree of deformation. It refers to the degree to which merozoites deform RBCs during invasion, and is given by a simplified four-point deformation scale (Weiss et al. 2015). We carried out a visual assessment of RBC deformation without prior knowledge of the Duffy RBC phenotype.

### Flickering spectroscopy

The biophysical properties of the RBCs were measured as previously described (Kariuki et al. 2020; Introini et al. 2022). Briefly, the Duffy RBC groups were each diluted to 0.01% HCT in incomplete RPMI 1640 to provide optimal cell density and avoid overlapping cells, and loaded into individual SecureSeal hybridisation chambers maintained at 37°C using the same microscope setup described above for video microscopy. We recorded 20s time-lapse videos at a high frame rate (514 fps) and a short exposure time (0.8 ms), to allow for the detection of RBC membrane contour. RBC contour was detected in bright-field for each frame with subpixel resolution by an optimised algorithm developed in-house by Cicuta Group at the Cavendish Laboratory and implemented in Matlab. The RBC equatorial contour was decomposed into fluctuation modes by Fourier transformation to give a fluctuation power spectrum of mean square mode amplitudes at the cell equator ⟨|*h*(*q_x_*, *y* = 0)|^2^⟩ as a function of mode wave vector (*qx*) (where *x* refers to the projection of modes *q* on the *x-*axis). From these data, RBC bending modulus (*κ*) and tension (*σ*) were fitted using the following equation:

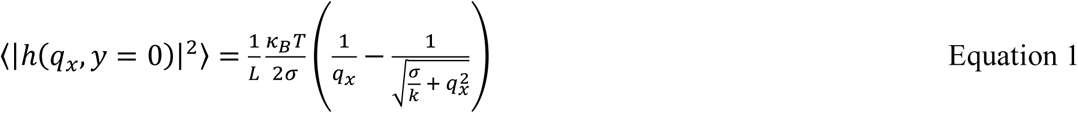

Where *k*_B_ is the Boltzmann constant, *T* is temperature, and *L* is the mean circumference of the RBC contour. The above equation considers fluctuation as having a thermal origin and is derived from the energy of deforming a flat sheet (Pécréaux et al. 2004), it is a good description of shape fluctuations of the cell’s equator in only a limited range of modes. We used modes of between 8 and 20 because lower modes fail due to the closed shape of a cell, whereas higher modes fail because of optical and temporal resolution. The RBC membrane fluctuations and their relaxation time also allowed for the quantification of RBC membrane viscosity (Introini 2020).

### RBC plasma membrane profiling

RBC plasma membrane profiling was performed as previously described (Ravenhill et al. 2019). Briefly, the 16 cryopreserved Duffy RBC pellets were thawed, and 200μl of each sample was washed twice in cold Phosphate-buffered saline (PBS) and resuspended in the remaining PBS between washes. The resuspended RBCs were further purified by Plasmodipur filtration, and RBC surface sialic acid residues were oxidised with sodium meta-periodate (Thermo Fisher) and biotinylated with aminooxy-biotin (Biotium). The reaction was quenched with glycerol and the RBCs were washed twice in cold PBS. After surface protein biotinylation, the intact RBCs were lysed with 1% Triton X-100 and biotinylated proteins were pulled down using high-affinity streptavidin agarose beads (Pierce) and washed extensively. The captured proteins on-beads were denatured with dithiothreitol, alkylated with iodoacetamide (Sigma), and digested with trypsin (Promega) in 200 mM HEPES (pH 8.5) for 3 hrs. Finally, each of the 16 peptide samples was separately labelled with a unique tandem mass tag (TMT; Thermo Fisher) reagents and quenched with 5% hydroxylamine for 15 mins. Equal volumes of the TMT-labelled peptide samples (10% of each sample) were combined in a single tube, and the mixture was subjected to C18 solid-phase extraction (SPE) using a custom-made stage tip before being dried with a vacuum concentrator and re-suspended in a mixture of 5% formic acid and 4% acetonitrile for mass spectrometry.

### Mass spectrometry and protein quantification

Mass spectrometry data were acquired using an Orbitrap Fusion Lumos (Thermo Scientific) interfaced via an EASYspray source to an Ultimate 3000 RSLC nano ultrahigh-performance liquid chromatography (UHPLC) column. Peptides were loaded onto an Acclaim PepMap nanoViper precolumn (internal diameter 100μm, height 2cm; Thermo Fisher) and resolved using a PepMap RSLC C18 EASYspray column (internal diameter 75μm, height 50cm, particle size 2μm). The loading solvent was 0.1% formic acid; the analytical solvent comprised solvents A (0.1% formic acid) and B (80% acetonitrile plus 0.1% formic acid). All separations were carried out at 40°C. Samples were loaded at 5μl min^-1^ for 5 min in loading solvent before beginning the analytical gradient. The following gradient was used: 3-7% solvent B over 2 min, 7-37% solvent B over 173 min, and a 4 min wash at 95% solvent B and equilibration at 3% solvent B for 15 min. Each analysis used a MultiNotch MS3-based TMT method (McAlister et al. 2014). The following settings were used: MS1, 380-1,500Th, 120,000 resolution, 2.0e^5^ automatic gain control (AGC) target, 50ms maximum injection time; MS2, quadrupole isolation at an isolation width of *m*/*z* 0.7, collision-induced dissociation (CID) fragmentation (normalised collision energy (NCE) 35) with ion-trap scanning in turbo mode with 1.5e^4^ AGC target, 120ms maximum injection time; MS3, in synchronous precursor selection mode the top ten MS2 ions were selected for HCD fragmentation (NCE 65) and scanned in the Orbitrap at a resolution of 60,000 with an AGC target of 1.0e^5^ and a maximum accumulation time of 150 ms. Ions were not accumulated for all parallelizable time. The entire MS/MS/MS cycle had a target time of 3s. Dynamic exclusion was set to ±10 p.p.m. for 70s. MS2 fragmentation was triggered on precursors of 5.0e^3^ counts and above. The data was processed using Proteome Discoverer v2.2 using the human Uniprot database (downloaded 13/01/21) with normalisation turned off. Oxidation (M) was set a potential variable modification, and carbamidomethyl (C) was set as a fixed modification.

As previously described (Ravenhill et al. 2019), the quantified proteins were filtered to include those most likely to be present with high confidence at the cell surface (Krogh et al. 2001). Contaminants, including serum, platelet, and leukocyte-related proteins, were removed before the proteins were normalised using the summed signal-to-noise values of the three most abundant proteins and less variable proteins (GYPA, GYPC, and Band 3). We used the method of significance A to estimate the *P* value that each protein ratio was significantly different to 1 (Cox and Mann 2008), and corrected for multiple comparisons using the Benjamini-Hochberg method.

### Equations of the evolutionary-epidemiological model

Equations 2-9 describe the rates of change of numbers of immature and mature susceptible (S_1_ and S_2_), potentially severely infected (V_1_ and V_2_), immune to severe malaria (R_1_ and R_2_) and mildly infected (I_1_ and I_2_) individuals of genotype *i*. For a more detailed discussion of a similar model, see (Penman and Gandon 2020).

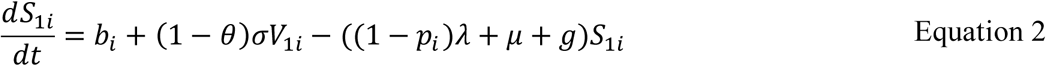

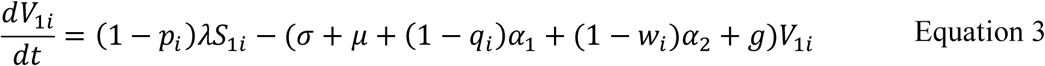

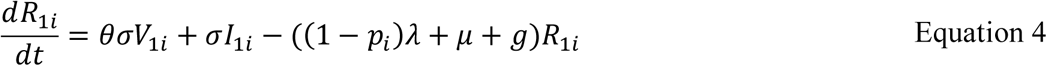

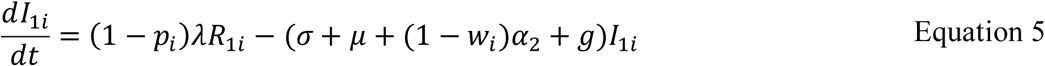

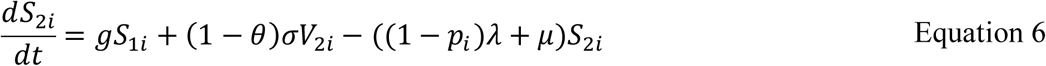

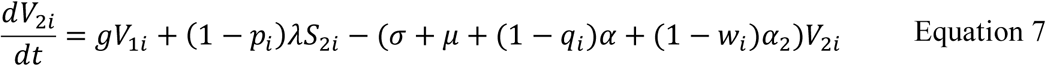

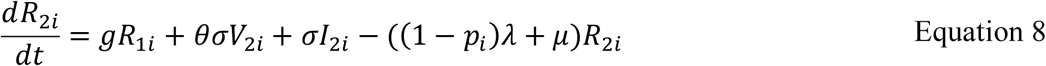

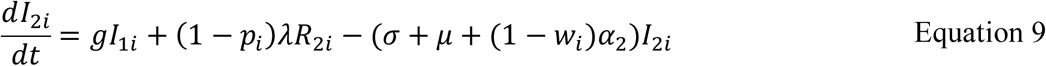

The force of infection, *λ*, is given by equation 10 where *N* = the total population size.

The birth rate, *b_i_* is given by equation 11, where *K*= the carrying capacity of the population; *r* = a fecundity parameter; *ψ* determines the reproductive cost of severe infection and *d_i_* determines the proportion of births which will be of genotype *i*. *d_i_* is calculated from the allele frequencies among the reproducing adult population according to the laws of Mendelian inheritance. Note that in the model which includes the sickle cell mutation, homozygotes for the sickle cell mutation are assumed to not survive, and thus (for simplicity) they do not enter the population at all; the proportions of births of each genotype are therefore rescaled to account for this loss.

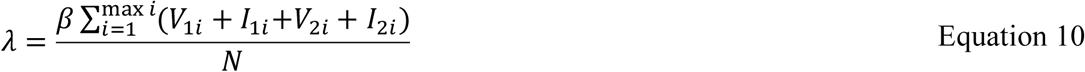

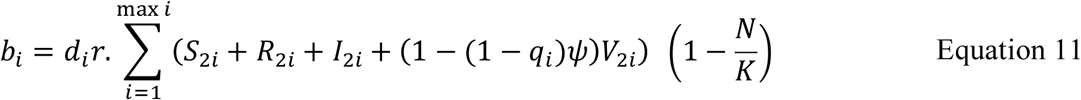

Parameter θ controls the proportion of hosts recovering out of compartment *V* who gain adaptive immunity, protecting them against death from severe malaria disease. The average number of infections experienced before protective immunity against severe disease is achieved is therefore equal to 1/ θ. Note that in this model, there is also a baseline malaria mortality rate (Table S7), not associated with severe malaria syndromes, which applies to all malaria infections. *p_i_* controls the protection against infection afforded to genotype *i*. If *p_i_* = 1, genotype *i* is fully protected against infection; if *p_i_* = 0, genotype *i* becomes infected at the maximum possible rate within the system. Parameter *q_i_* controls the protection against severe malaria syndromes afforded to genotype *i*. If *q_i_* = 1, genotype *i* is fully protected against severe malaria mortality and against the reproductive costs of virulent malaria. Parameter *w_i_* controls the protection against baseline malaria mortality afforded to genotype *i*. If *w_i_* = 1, genotype *i* is fully protected against baseline malaria mortality. All parameters of the model, and the values used here, are given in Table S7.

