## Supplementary Information for "Invasion preferences suggest a possible role for *Plasmodium falciparum* parasites in the expansion of Duffy negativity in West and Central Africa"

### Supplementary Figures

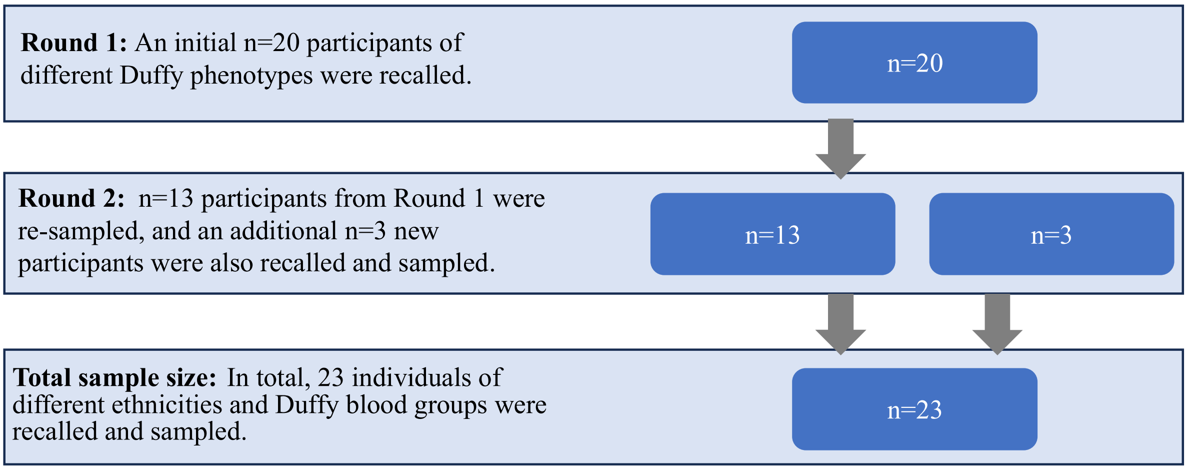


**Figure S1: Sample collection and study design.** The Duffy blood samples were collected in two recall-by-genotype study rounds in 2021 (round 1) and 2022 (round 2), from 20 participants in round 1 and 16 participants in round 2 (13 of the 20 participants from round 1 were re-sampled, and three new participants were added). The round 1 samples were received in batches of five (i.e., five bi-weekly batches and five samples per batch), and only three of the four major Duffy blood groups were represented in each batch, while in round 2, samples were collected in four bi-weekly batches (four samples per batch) and all four Duffy phenotypes were represented in each of the four batches. The presence of all four Duffy blood groups in a batch was useful for the preference assays as it meant that RBCs from all four samples could be labelled with different fluorescent dyes and incubated together with malaria parasites in the same well, allowing for the comparison of invasion into all four different RBC samples simultaneously.

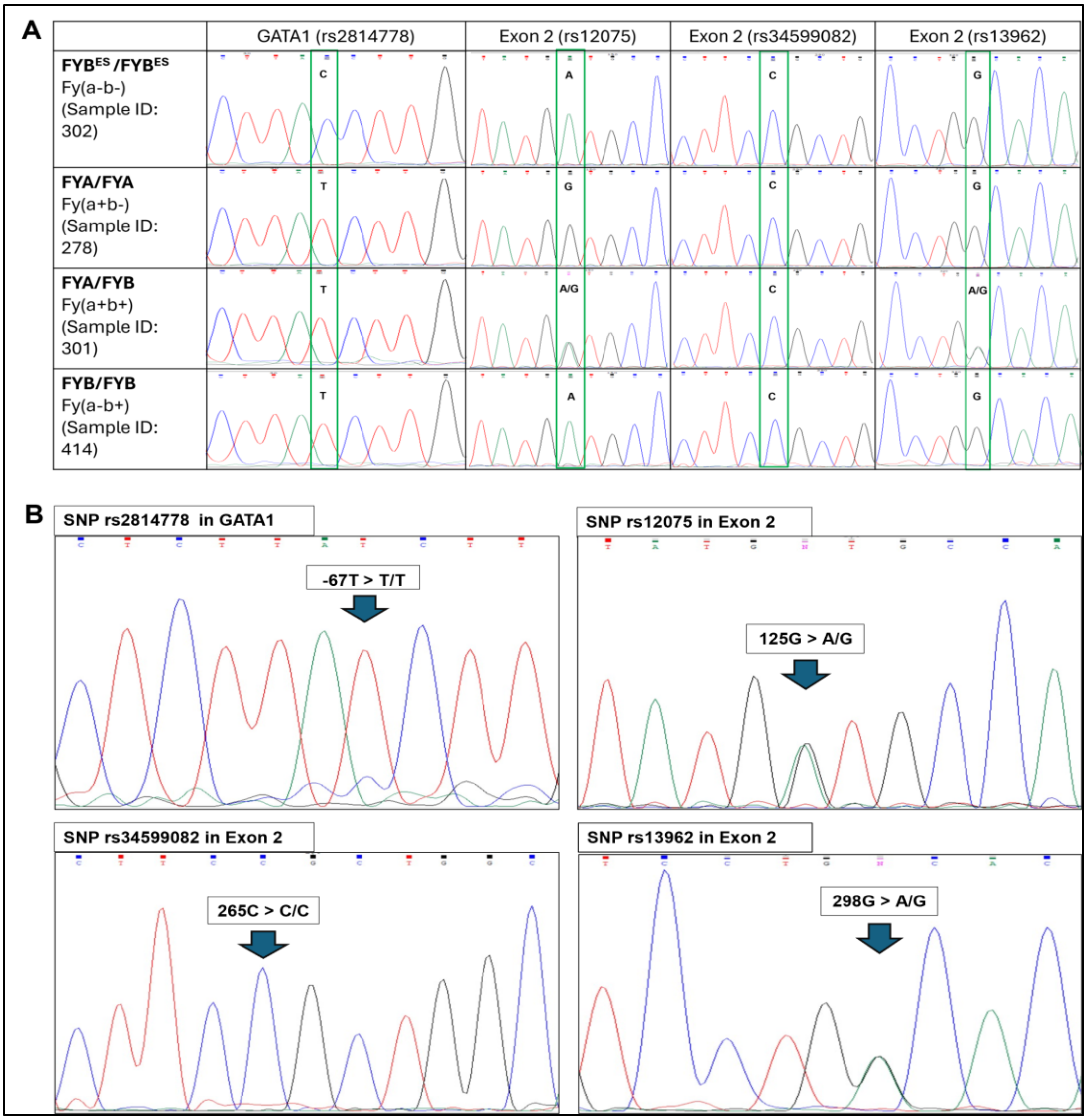


**Figure S2: Segments of chromatograms of Duffy-positive and -negative samples.**
(**A**) Chromatogram of four Duffy samples showing the GATA1 mutation (-67 T>C; rs2814778) for a Duffy-negative sample (sample 302), and the wildtype SNP (-67 T>T; rs2814778) for homozygous FYA sample (FYA/FYA; sample 278), heterozygous (FYA/FYB; sample 301), and homozygous FYB sample (FYB/FYB; sample 414). The positions of all SNPs both in the GATA1 (rs2814778) and Exon 2 (rs12075; rs34599082; and rs13962) are highlighted in green rectangles for all samples.
(**B**) Segments of a chromatogram from a Duffy-positive sample (sample 559) in round 2 that was genotyped as FYA/FYB (heterozygous) showing SNP locations in the GATA1 and Exon 2 regions of the gene. The agglutination test showed that the donor has the Fy(a+b-) phenotype; however, the sequencing results above showed that the individual has no mutation in GATA1 (-67 T> T/T; wildtype). The SNP location (125G>A/G; rs12075) determines the presence of the FYA or FYB alleles and their corresponding phenotypes. Two peaks of similar intensity are seen at position 125 in Exon 2, meaning the individual has the co-dominant FYA and the FYB alleles at this locus (i.e., 125G > A/G) and would express both $Fy^a/Fy^b$ antigens and the phenotype Fy(a+b+). No mutation is seen at position 265, the third SNP location (rs34599082). A mutation is present at position 298 in Exon 2, and two peaks of relatively similar intensities are seen, indicating that the individual has both the FYA and FYB alleles at this locus, and the donor was assigned the Fy(a+b+) phenotype.

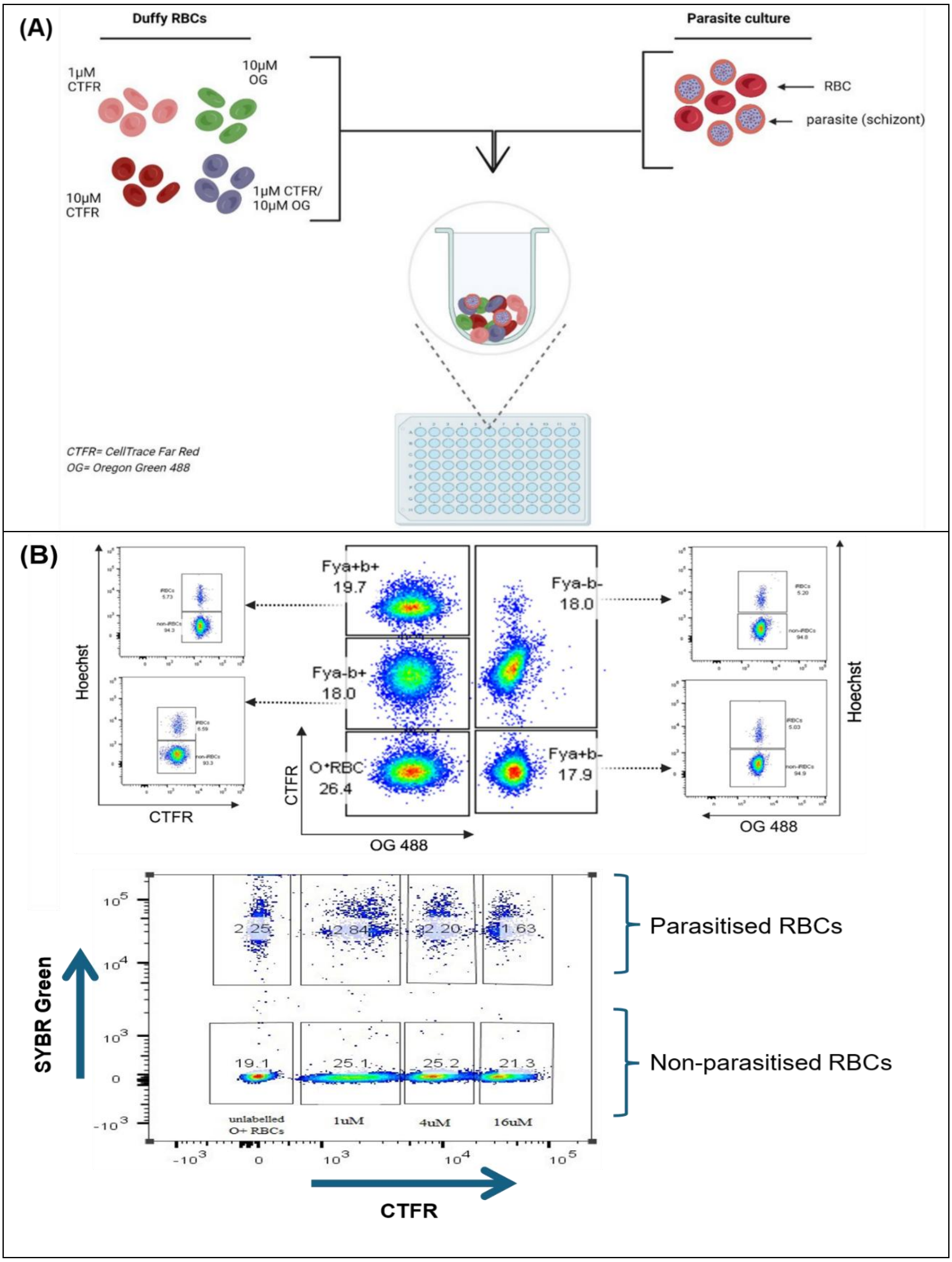
(A)
Duffy RBCs
Parasite culture
1µM CTFR
10µM OG
10µM CTFR
1µM CTFR/ 10µM OG
RBC
parasite (schizont)
CTFR= CellTrace Far Red
OG= Oregon Green 488
(B)
Fya+b+ 19.7
Fya-b+ 18.0
O+RBC 26.4
Fya-b- 18.0
Fya+b- 17.9
Hoechst
CTFR
OG 488
SYBR Green
2.25
2.84
2.20
1.63
19.1
25.1
25.2
21.3
unlabelled O+ RBCs
1uM
4uM
16uM
Parasitised RBCs
Non-parasitised RBCs

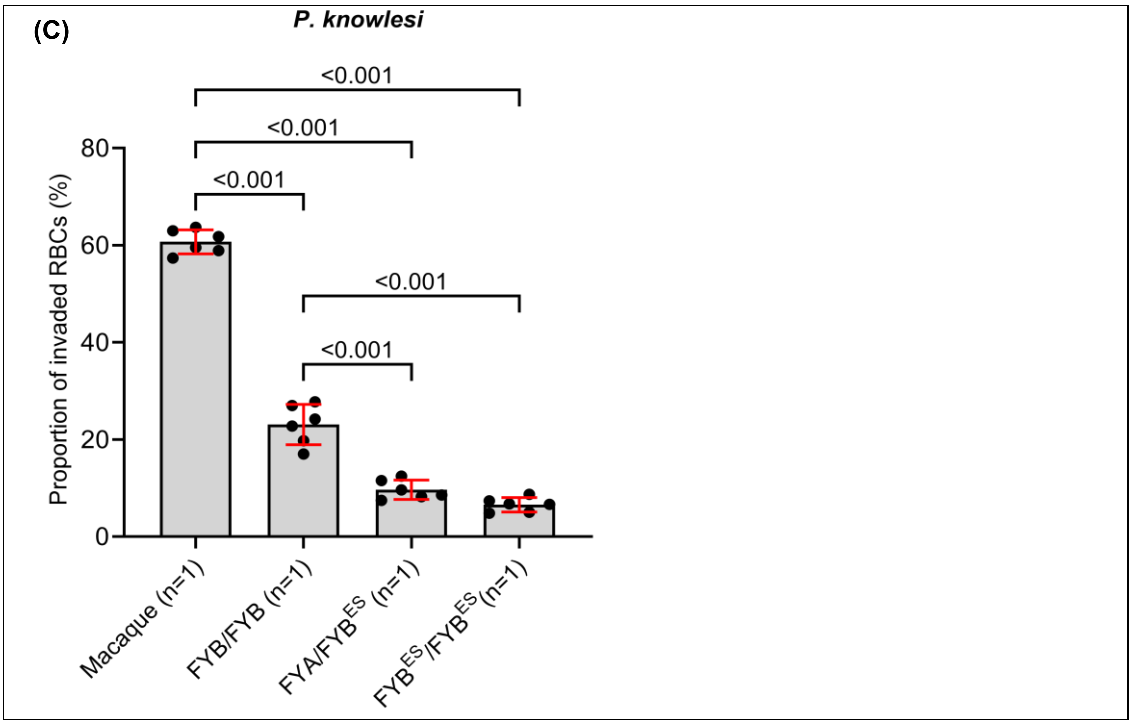


**Figure S3: The *in vitro* parasite preference assay.**
(**A**) Schematic of the *in vitro* preference assay based on an earlier version (Theron et al. 2018).
(**B**) Top panel - gating strategy used for the round 2 samples (RBCs from all 4 Duffy phenotypes were incubated together with parasites in a single well), showing the proportion of invaded RBCs by *P. falciparum* in a single well of a preference assay after debris and doublets were excluded. Each of the Duffy RBCs contained both parasitised and non-parasitised RBCs and was gated using the fluorescent intensities of CTFR vs Hoechst (to quantify CTFR-labelled parasitised RBCs) and OG vs Hoechst (to quantify OG-labelled parasitised RBCs). Bottom panel -gating strategy used for round 1 samples, showing the parasitised and non-parasitised RBCs after debris and doublets were excluded. The RBCs were labelled with increasing concentrations of CTFR, and the percentage of parasitised RBCs in each labelled RBC group was determined using the fluorescence intensity of CTFR vs SYBR Green dyes.
(**C**) We assessed the sensitivity of the above preference assay using Macaque and human RBCs prior to analysing the Duffy samples. The macaque RBCs (of unknown genotype) were preferentially invaded over the human RBCs, and among the human RBCs, the FYB/FYB RBCs were preferentially invaded over the other genotypes. On average, 60% (±2.5) of the Macaque RBCs were invaded compared to 23.1% (±4.2) for FYB/FYB, 9.7% (±2.0) for FYA/FYB$^{ES}$, and 6.6% (±1.5) for FYB$^{ES}$/FYB$^{ES}$ RBCs. These data clearly show that the preference assay is highly sensitive and able to reliably detect even minor differences in the invasion preference of human malaria parasites. The assay was performed as described in the Methods, using the same *P. knowlesi* parasites and human RBCs, and each assay was performed in triplicate, resulting in six replicates for each donor sample. Statistical comparisons between the samples were performed using a one-way ANOVA, and a Tukey's test was used to correct for multiple comparisons. The mean, standard deviation, and statistically significant differences are shown for each sample.

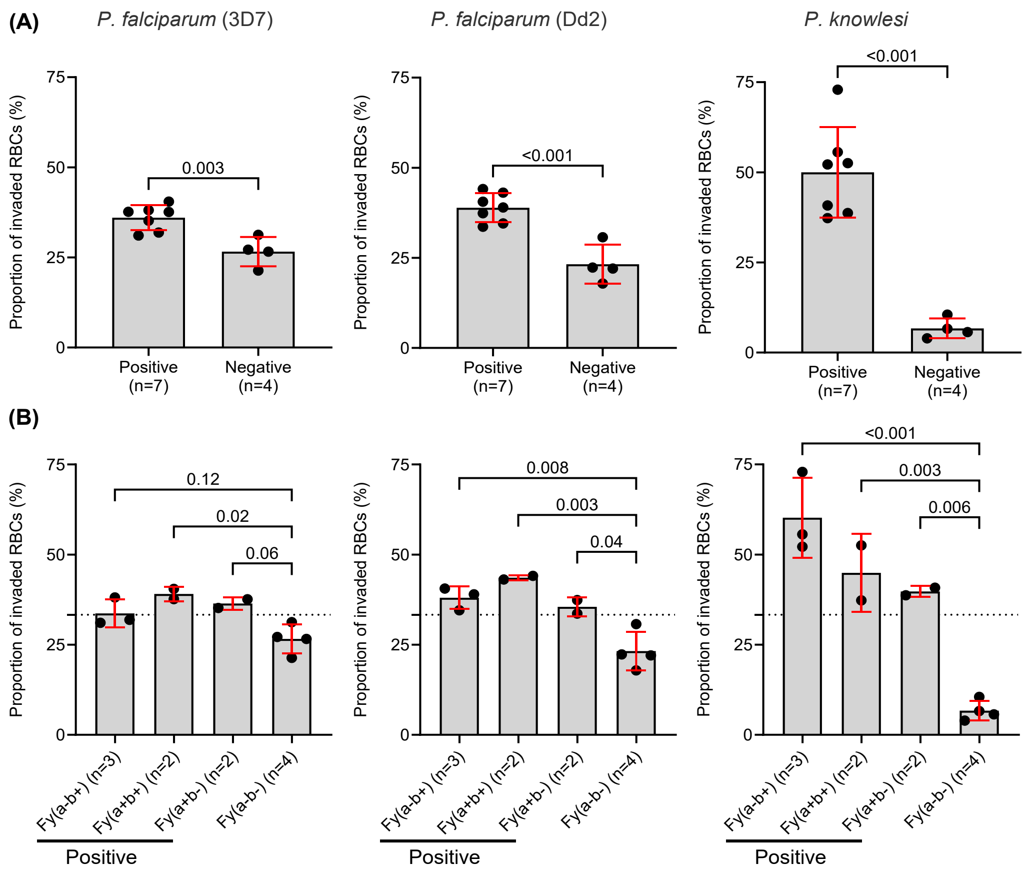


**Figure S4: The invasion preference of malaria parasites for Duffy RBC groups in round 1.** In round 1, only three Duffy phenotypes were compared at a time in the preference assay, as shown in Figure S3B, and all parasites showed a clear preference for RBCs from Duffy-positive individuals.

(**A**) The preference of *P. falciparum* parasites was skewed towards Duffy-positive RBCs and away from Duffy-negative RBCs. When grouped by Duffy status (i.e., Duffy-positive vs Duffy-negative), the average proportion of invaded RBCs by 3D7 parasites was 36.0% (±3.5) for the Duffy-positive and 26.6% (±4.1) for the Duffy-negative RBCs, respectively. The average invasion for Dd2 parasites was 38.9% (±4.0) for Duffy-positive and 23.2% (±5.4) for Duffy-negative RBCs. As expected, invasion of the Duffy-negative RBCs by *P. knowlesi* was significantly reduced relative to Duffy-positive RBCs; the average proportion of invaded RBCs was 50.0% (±12.5) for Duffy-positive compared to 6.7% (±2.8) for Duffy-negative RBCs.

(**B**) As several Duffy genotypes/phenotypes are represented in the Duffy-positive samples, we re-analysed the data to compare invasion rates in all four Duffy phenotypes. The average proportion of invaded RBCs with 3D7 was 33.7% (±3.9), 39.1% (±2.0), 36.4% (±1.7), and 26.6% (±4.1) for Fy(a-b+), Fy(a+b+), Fy(a+b-), and Fy(a-b-) RBCs, respectively. The Dd2 parasites also showed an invasion preference for Duffy-positive RBCs over Duffy-negative RBCs with an average invasion of 38.0% (±3.1), 43.6% (±0.7), 35.5% (±2.7) for Fy(a-b+), Fy(a+b+), and Fy(a+b-) RBCs, respectively, compared to 23.2% (±5.4) for the Duffy-negative RBCs. The average proportion of invaded RBCs for the *P. knowlesi* parasites was 60.2% (±11.1), 44.9% (±10.8), and 39.8% (±1.5) for the Fy(a-b+), Fy(a+b+), and Fy(a+b-) RBCs, respectively, compared to 6.7% (±2.8) for the Duffy-negative RBCs. Statistical comparison for data in (A) was carried out using a two-tailed t-test, and comparisons between the four RBC groups in (B) were carried out using One-way ANOVA, followed by pairwise comparisons using Tukey's HSD (honestly significant difference) test. The mean and standard deviation are shown for each group.

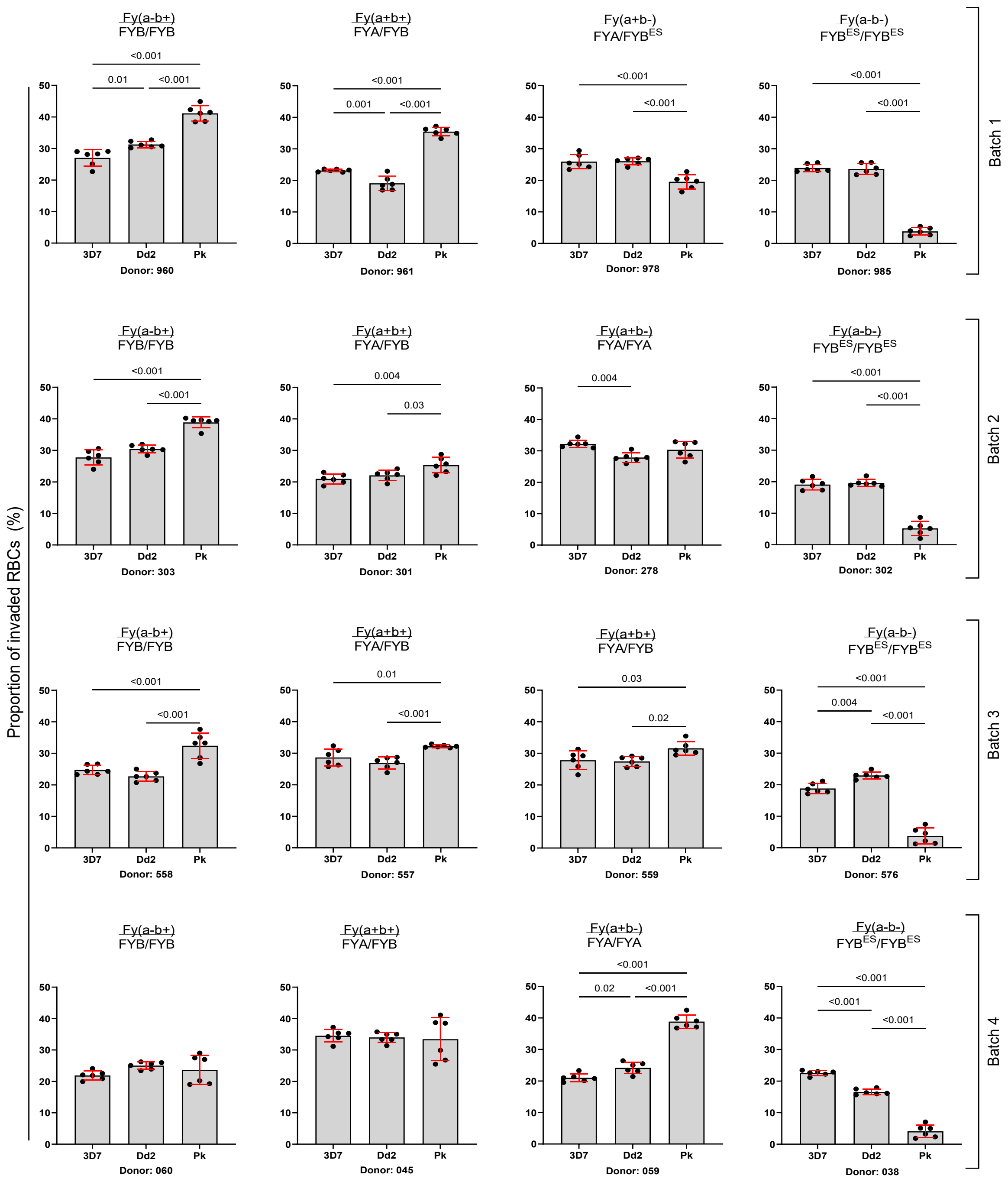


**Figure S5: The invasion rates of three malaria parasites in donor RBC sample tested in 2.** Each donor RBC sample was tested with *P. falciparum* (3D7 and Dd2 strains) and *P. knowlesi* (Pk) parasites in triplicate, and each assay was duplicated to produce six replicates per donor; all six replicates are shown. The corresponding Duffy phenotype and genotype are listed above each sample. In Batch 3, the agglutination test showed that donor 559 has the Fy(a+b-) phenotype, however, the genotyping showed that the donor is FYA/FYB (heterozygous) (Figure S4B). The bar plots are coloured based on Duffy RBC phenotype (blood group), and the *y-axis* represents the proportion of invaded RBCs in each sample by the tested parasites. Statistical comparisons between the groups were carried out using One-way ANOVA, followed by pairwise comparisons using Tukey's test. The mean and standard deviations are shown for each parasite strain/species, and all statistically significant *P* values (i.e. $P \leq 0.05$) are shown.

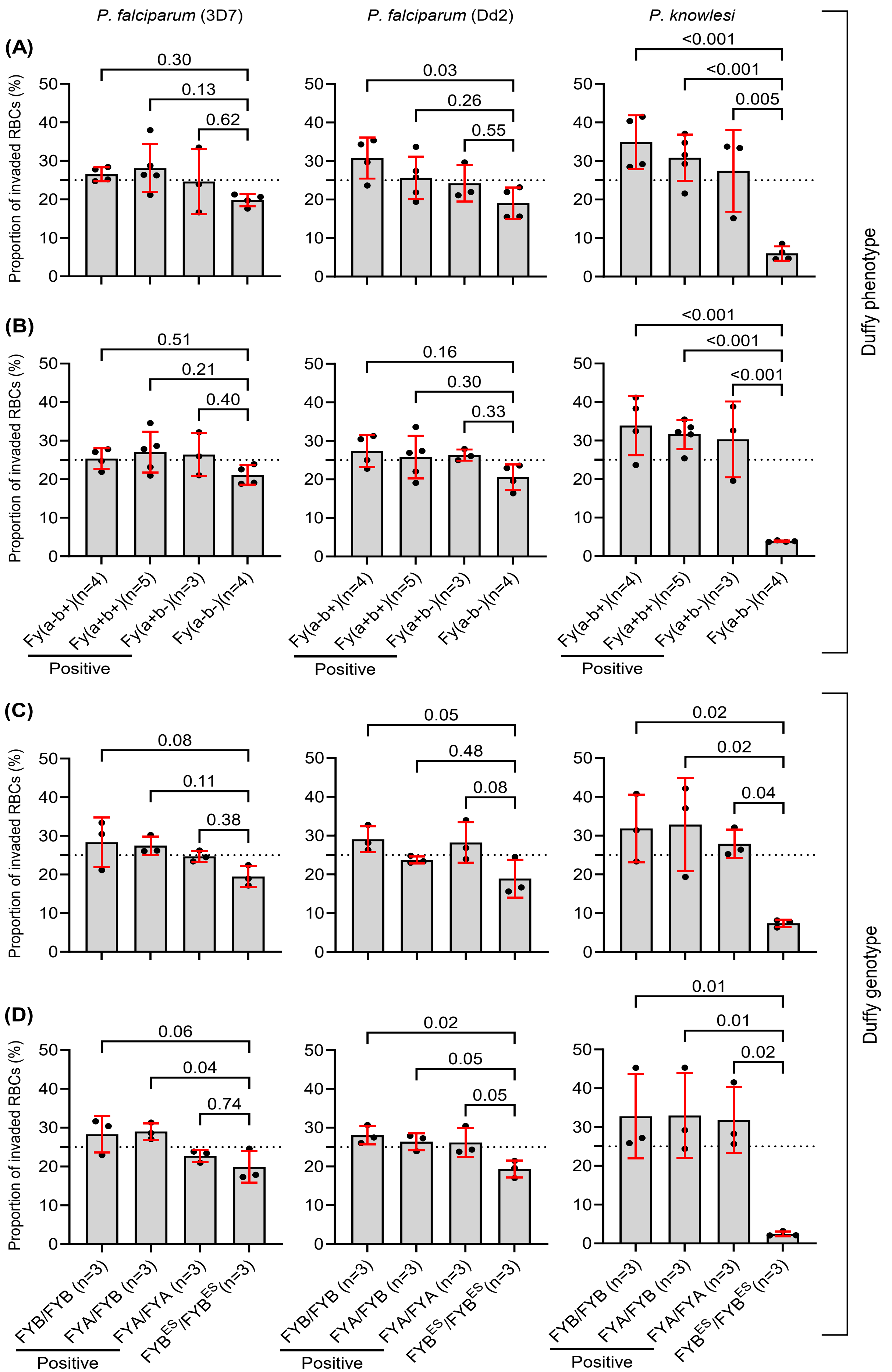
P. falciparum (3D7)
P. falciparum (Dd2)
P. knowlesi
(A)
(B)
(C)
(D)
Proportion of invaded RBCs (%)
0.30
0.13
0.62
0.03
0.26
0.55
<0.001
<0.001
0.005
0.51
0.21
0.40
0.16
0.30
0.33
<0.001
<0.001
<0.001
Fy(a-b+)(n=4)
Fy(a+b+)(n=5)
Fy(a+b-)(n=3)
Fy(a-b-)(n=4)
Positive
Duffy phenotype
0.08
0.11
0.38
0.05
0.48
0.08
0.02
0.02
0.04
0.06
0.04
0.74
0.02
0.05
0.05
0.01
0.01
0.02
FYB/FYB (n=3)
FYA/FYB (n=3)
FYA/FYA (n=3)
FYB^ES/FYB^ES (n=3)
Positive
Duffy genotype

**Figure S6: Effect of Duffy phenotype and genotype on RBC invasion by *P. falciparum* and *P. knowlesi* parasites.**
(**A**) The Duffy phenotypes of the grouped data presented in Figure 1A and C.
(**B**) The Duffy phenotypes of the grouped data presented in Figure 1B and D.
(**C**) To assess the impact of Duffy genotype on the invasion preference of the malaria parasites, we performed additional sets of preference assays using the same RBC samples and parasites as in Figure 1 and Figure S4, to compare invasion only by genotype rather than phenotype.
(**D**) The Duffy genotypes were also tested in invasion assays as in (C) above. All six replicates were averaged for each donor and shown on the plot. Statistical comparisons were performed using a one-way ANOVA, followed by Tukey's test.

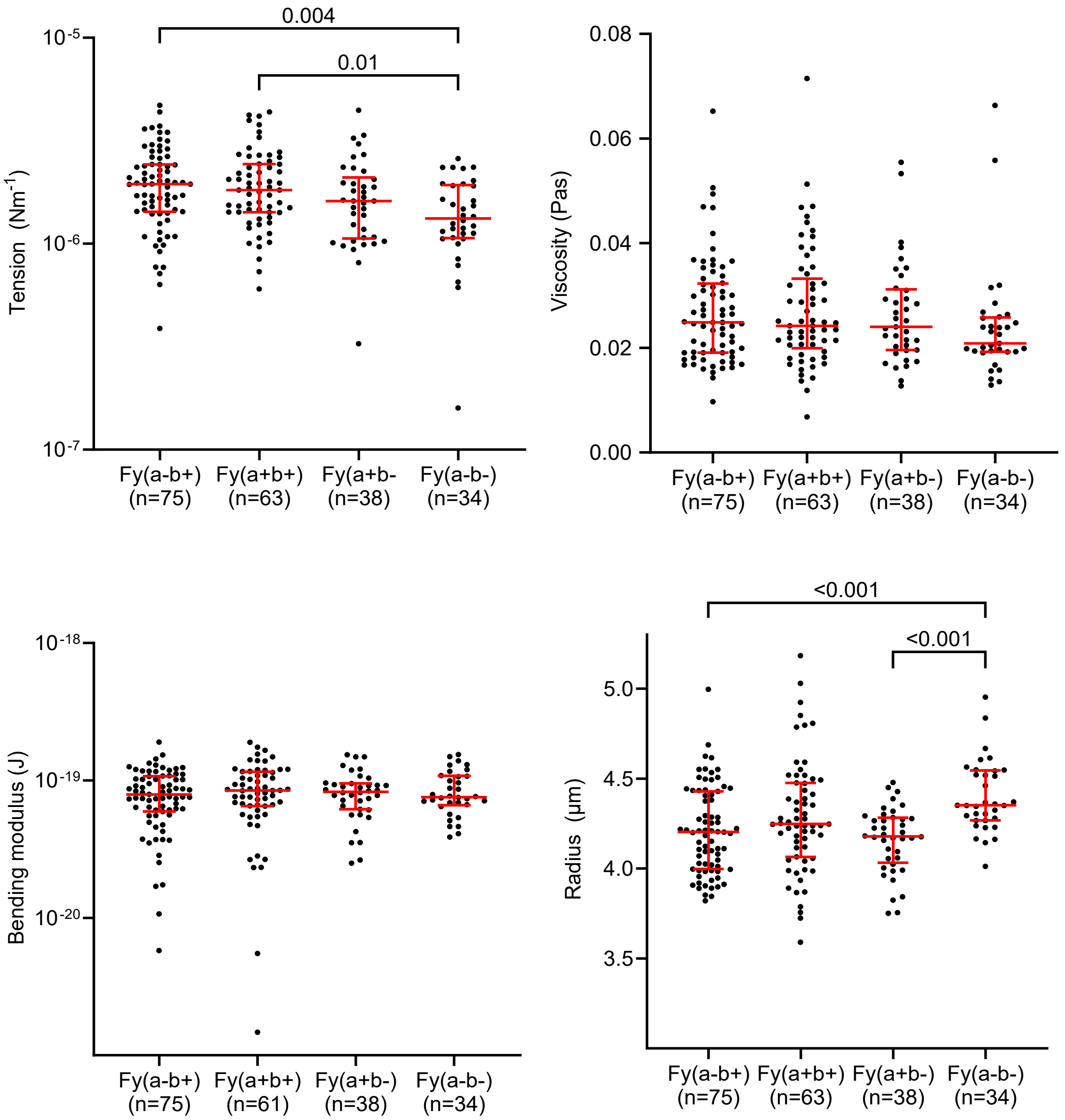


**Figure S7: Flickering analysis of Duffy RBC samples collected in round 2.** The Duffy RBC samples analysed in this study were collected in two rounds, as summarised in **Figure S1.** As all samples except three (3) samples were initially collected in round 1 and analysed, only a randomly selected subset of the samples (n=6) was therefore analysed in round 2 – (Fy(a-b+)(n=2), Fy(a+b+)(n=2), Fy(a+b-)(n=1), and Fy(a-b-)(n=1). Statistical significance between the four phenotypes was determined by a Kruskal-Wallis test, and differences were corrected for multiple comparisons using Dunn's test. The median, IQR, and statistically significant differences are shown for each phenotype.

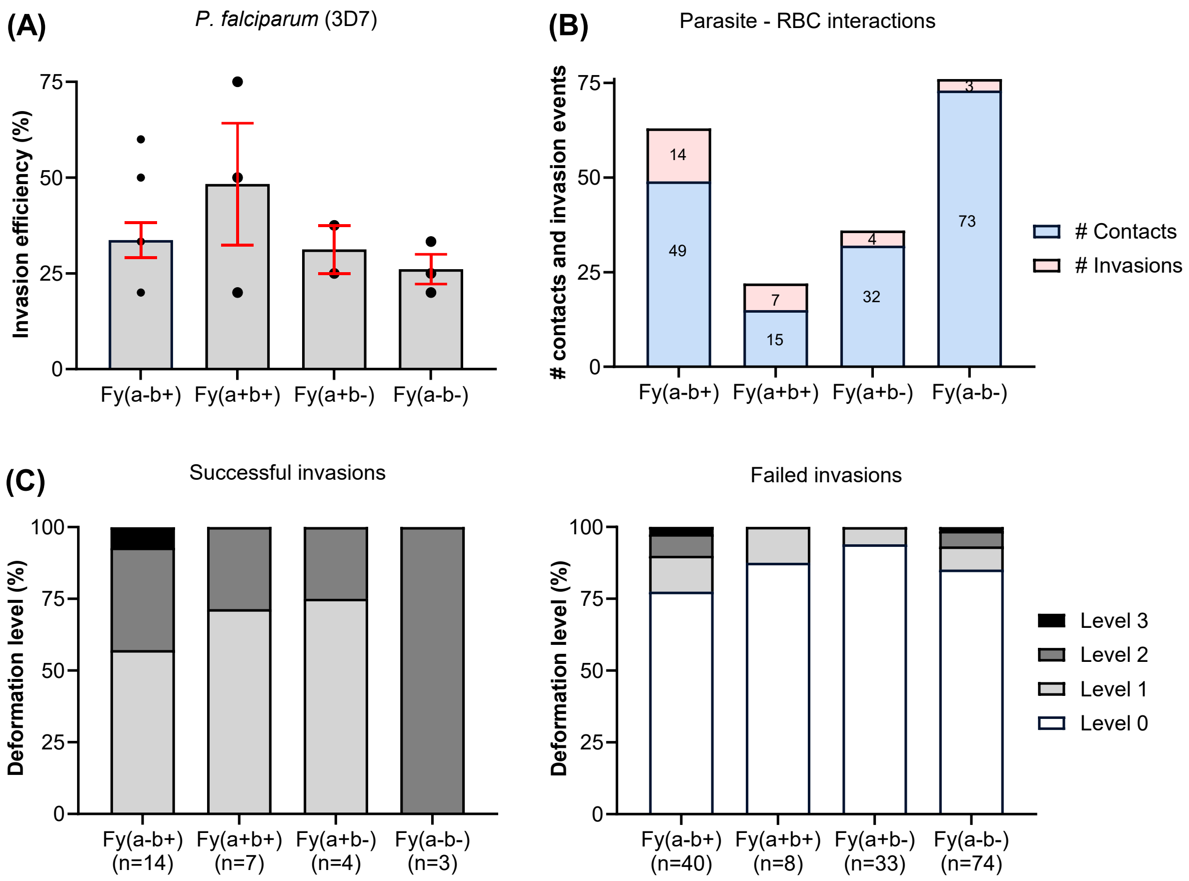


**Figure S8: Invasion efficiency and RBC deformation by 3D7 parasites.**
(**A**) The invasion efficiency of 3D7 parasites was quantified by video microscopy. The following RBC groups were analysed: Fy(a-b+)(n=3); Fy(a+b+)(n=2); Fy(a+b-)(n=2); and Fy(a-b-)(n=3). The results showed no statistically significant differences in invasion efficiency between the RBC samples. Statistical comparisons were carried out using One-way ANOVA, and Tukey's test was used to determine the significant differences between the groups. The bar plot shows the mean and standard error of the mean (SEM) for each Duffy RBC group.
(**B**) The number of times merozoites came into contact with RBCs and successfully invaded is shown for each Duffy blood group. The ratio of successful invasions and the number of merozoites that came in contact with RBCs was used to determine the invasion efficiency in (A) above.
(**C**) The deformation levels of successful and failed invasion events were also determined. The number below each phenotype 'n' represents the number of merozoites that deformed the RBCs and either failed or successfully invaded each blood group.

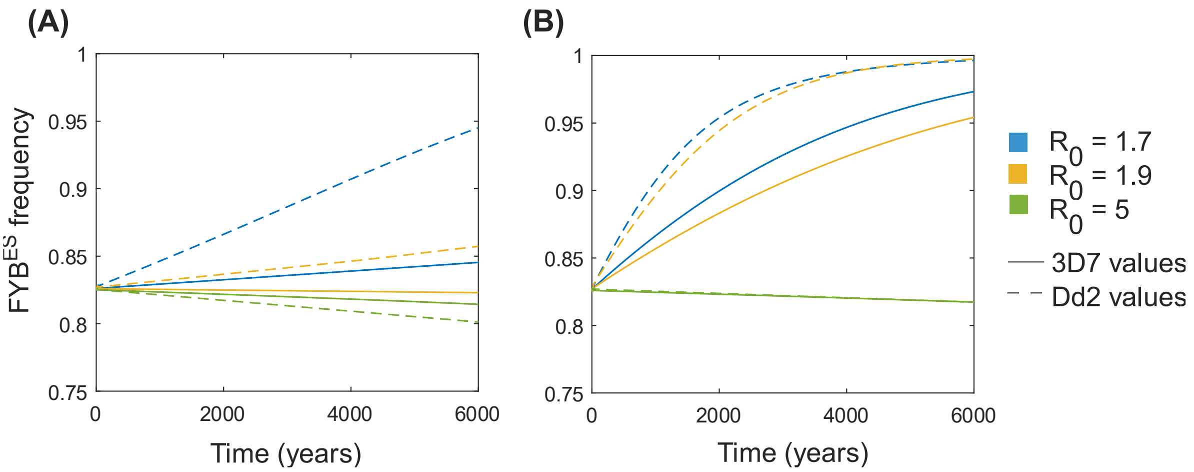


**Figure S9: Sensitivity analysis of Figure 4.** Figures (A) and (B) both use a starting frequency of 0.825 $FYB^{ES}$. In panel (A) $\theta = 1/5$ , $\alpha_1$=0.04 and $\alpha_2$= 0.00005 . In (B), $\theta = 1/20$ , $\alpha_1$= 0.005 and $\alpha_2$= 0.00005. The relationship that we predict (i.e. that *P. falciparum* is most likely to elevate $FYB^{ES}$ frequencies at relatively low levels of transmission) is not dependent on the exact parameter values that were used in Table S7. To illustrate this, Figure S9 re-generates two new versions of Figure 4A, but using different malaria mortality rates and assuming that immunity to *P. falciparum* virulence is gained even more rapidly than in Figure 4. The rates of allele frequency change are different, but the pattern that $FYB^{ES}$ frequencies increase more rapidly with lower *P. falciparum* transmission remains.

### Supplementary Tables

**Table S1: Demographic characteristics and sample parameters of study participants.**

| **Round 1** | | | | | |
|---|---|---|---|---|---|
| **Characteristics** | **Fy(a-b+)** | **Fy(a+b+)** | **Fy(a+b-)** | **Fy(a-b-)** | **Fy(a+b$^{w}$)** |
| Sample size (n) | 6 | 5 | 3 | 5 | 1 |
| Age in years (sd) | 44 (13.78) | 27.4 (5.13) | 30.7 (7.51) | 37 (15.15) | 24 |
| Sex (M/F) | 3/3 | 3/2 | 2/1 | 3/2 | 1/0 |
| White blood cell count ($10^9$ L) (sd) | 6.25 (2.51) | 7.99 (2.70) | 4.91 (0.66) | 5.11 (0.96) | 8.34 |
| Red blood cell count ($10^{12}$ L) (sd) | 4.73 (0.39) | 5.01 (0.36) | 4.84 (0.34) | 4.64 (0.47) | 6.0 |
| Haemoglobin (g/L) (sd) | 142.7 (8.02) | 137.8 (16.54) | 147 (8.0) | 129.6 (13.22) | 117 |
| Haematocrit (L/L) (sd) | 0.429 (0.023) | 0.426 (0.038) | 0.438 (0.019) | 0.391 (0.042) | 0.359 |
| Mean cell volume (fL) (sd) | 91.15 (4.05) | 85 (2.67) | 90.2 (4.35) | 84.48 (7.59) | 59.8 |
| Mean cell haemoglobin (pg) (sd) | 30.23 (1.46) | 27.44 (1.65) | 30.27 (1.54) | 28.06 (3.21) | 19.5 |
| Mean cell haemoglobin concentration (g/L) (sd) | 332 (6.07) | 322.6 (11.48) | 335.7 (3.06) | 332 (17.35) | 326 |
| Platelet ($10^9$ L) (sd) | 267 (111.8) | 265 (55.65) | 194 (8.96) | 195 (58.48) | 196 |
| RBC distribution width (%) (sd) | 12.85 (0.75) | 13.66 (1.42) | 12 (0.44) | 13.84 (2.56) | 18.5 |
| **Round 2** | | | | | |
| Sample size (n) | 4 | 5 | 3 | 4 | - |
| Age in years (sd) | 46.5 (9.95) | 27.2 (4.09) | 31.33 (8.08) | 30.5 (13.03) | - |
| Sex (M/F) | 1/3 | 4/1 | 2/1 | 3/1 | - |
| White blood cell count ($10^9$ L) (sd) | 6.91 (2.38) | 6.58 (2.32) | 5.08 (0.77) | 4.66 (0.89) | - |
| Red blood cell count ($10^{12}$ L) (sd) | 4.69 (0.29) | 5.04 (0.37) | 4.84 (0.42) | 5.14 (0.46) | - |
| Haemoglobin (g/L) (sd) | 139.00 (4.24) | 141.60 (13.26) | 144.67 (6.81) | 136.50 (11.21) | - |
| Haematocrit (L/L) (sd) | 0.42 (0.01) | 0.43 (0.03) | 0.44 (0.02) | 0.43 (0.02) | - |
| Mean cell volume (fL) (sd) | 90.55 (4.23) | 86.12 (3.18) | 91.17 (3.37) | 83.88 (7.31) | - |
| Mean cell haemoglobin (pg) (sd) | 29.70 (1.16) | 28.14 (1.37) | 30.00 (1.39) | 26.68 (3.01) | - |
| Mean cell haemoglobin concentration (g/L) (sd) | 328.25 (4.35) | 326.40 (8.41) | 328.67 (3.21) | 317.75 (9.91) | - |
| Platelet ($10^9$ L) (sd) | 267 (116.72) | 235.20 (52.10) | 196.67 (21.36) | 205.00 (56.02) | - |
| RBC distribution width (%) (sd) | 13.05 (0.42) | 12.92 (1.27) | 12.03 (0.55) | 14.50 (3.04) | - |

The table shows the demographic characteristics and blood sample parameters of study participants from both round 1 and round 2 at the time of sample collection. The mean values and standard deviations (sd) are shown for the age variable and RBC indices for each Duffy blood group. The sample size for each Duffy blood group is represented by the letter n in parenthesis (n); for the sex variable, F = female and M = male; and W = weak expression of the Fyb antigen on RBCs.

**Table S2: Duffy genotypes and phenotypes of study participants in both rounds 1 and 2.**

| Sample ID | Ethnicity | ABO | ABCA7 | Genotype | Phenotype | Fy Antigen | Year sampled |
|---|---|---|---|---|---|---|---|
| 950/960 | Asian (South) | A | Wildtype | FYB/FYB | Fy(a-b+) | Fyb+ | 2021/2022 |
| 800/961 | White | B | Wildtype | FYA/FYB | Fy(a+b+) | Fya+Fyb+ | 2021/2022 |
| 798/978 | Mixed | B | Wildtype | FYA/FYB$^{ES}$ | Fy(a+b-) | Fya+ | 2021/2022 |
| 412/985 | Black | O | Wildtype | FYB$^{ES}$/FYB$^{ES}$ | Fy(a-b-) | No antigen | 2021/2022 |
| 949/278 | White | O | Wildtype | FYA/FYA | Fy(a+b-) | Fya+ | 2021/2022 |
| 796/301 | Mixed | A | Wildtype | FYA/FYB | Fy(a+b+) | Fya+Fyb+ | 2021/2022 |
| 297/302 | Black | A | Wildtype | FYB$^{ES}$/FYB$^{ES}$ | Fy(a-b-) | No antigen | 2021/2022 |
| 947/303 | White | O | Wildtype | FYB/FYB* | Fy(a-b+) | Fyb+ | 2021/2022 |
| 416/557 | White | A | Wildtype | FYA/FYB | Fy(a+b+) | Fya+Fyb+ | 2021/2022 |
| 415/558 | White | O | Wildtype | FYB/FYB | Fy(a-b+) | Fyb+ | 2021/2022 |
| 559 | White | A | Wildtype | FYA/FYB | Fy(a+b+) | Fya+b+ | 2022 |
| **300/576** | **Black** | **O** | **Het** | **FYB$^{ES}$/FYB$^{ES}$** | **Fy(a-b-)** | **No antigen** | **2021/2022** |
| **038** | **Black** | **B** | **Het** | **FYB$^{ES}$/FYB$^{ES}$** | **Fy(a-b-)** | **No antigen** | **2022** |
| 299/045 | White | A | Wildtype | FYA/FYB | Fy(a+b+) | Fya+Fyb+ | 2021/2022 |
| 797/059 | Asian (East) | O | Wildtype | FYA/FYA | Fy(a+b-) | Fya+ | 2021/2022 |
| 060 | Black | O | Wildtype | FYB/FYB$^{ES}$ | Fy(a-b+) | Fyb+ | 2022 |
| 799 | Black | O | Hom | FYB$^{ES}$/FYB$^{ES}$ | Fy(a-b-) | No antigen | 2021 |
| 413 | Asian (South) | AB | Wildtype | FYA/FYX2 | Fy(a+b$^{weak}$) | Fya+Fyb$^{weak}$ | 2021 |
| 414 | Other | O | Wildtype | FYB/FYB | Fy(a-b+) | Fyb+ | 2021 |
| 298 | Mixed | O | Wildtype | FYA/FYB | Fy(a+b+) | Fya+Fyb+ | 2021 |
| 301 | White | A | Wildtype | FYB/FYB | Fy(a-b+) | Fyb+ | 2021 |
| 946 | Black | O | Het | FYB$^{ES}$/FYB$^{ES}$ | Fy(a-b-) | No antigen | 2021 |
| 948 | White | O | Wildtype | FYB/FYB | Fy(a-b+) | Fyb+ | 2021 |

1. A total of 20 donors were sampled in 2021 (round 1), and 13 of them were re-sampled again in 2022 (round 2). Three new donors (559, 038, and 060) were added in 2022 to make up for the 2021 donors that were not available in 2022.
2. Sample ID column represent the last 3 digits of the sample IDs for participants sampled in either 2021, 2022 or both years. For example, participants sampled/recalled in both 2021 and 2022 were give a unique sample ID for each year (round) which is separated by "/".
3. In the rare FYX2 allele, the presence of the 298G>A and 265C>T can decrease the expression of Fyb+ antigen resulting in a weak phenotype (Olsson et al. 1998), as is the case with sample 413, highlighted in blue above.
4. All ethnicities are self-reported; and *ES* = erythrocyte silent (i.e., lack of Fy antigen expression only on RBCs)
5. The expression of ABCA7 was downregulated in the Duffy-negative RBC samples (Figure 3), thus all samples were genotyped for a 44bp deletion (rs142076058) that disrupt the expression of the protein, we found three (3) heterozygous (3003/576, 038, 946) and one (1) homozygous samples with the deletion, however, the deletions does not correlate with protein expression as the relative abundance of ABCA7 was also decreased in samples that are wildtype for the deletion, as shown in Figure 3A.

**Table S3: Distribution of Duffy genotypes and phenotypes in our study population**

| Genotype | Phenotype | Asian | Black | Caucasian | Mixed/ Other | Number of Samples |
|---|---|---|---|---|---|---|
| FYA/FYA | Fy(a+b-) | 1 | 0 | 1 | 0 | 2 |
| FYB/FYB | Fy(a-b+) | 1 | 0 | 4 | 1 | 6 |
| FYA/FYB | Fy(a+b+) | 0 | 0 | 4 | 2 | 6 |
| FYA/FYB$^{ES}$ | Fy(a+b-) | 0 | 0 | 0 | 1 | 1 |
| FYB/FYB$^{ES}$ | Fy(a-b+) | 0 | 1 | 0 | 0 | 1 |
| FYB$^{ES}$/FYB$^{ES}$ | Fy(a-b-) | 0 | 6 | 0 | 0 | 6 |
| FYA/FYX2 | Fy(a+b$^{weak}$) | 1 | 0 | 0 | 0 | 1 |
| | **Total** | 3 | 7 | 9 | 4 | 23 |

### Table S4. Differentially expressed proteins between Duffy-positive and -negative RBCs

| Accession | Gene | Protein name | Peptides | *P* value | Relative abundance in Duffy-negative |
|---|---|---|---|---|---|
| Q16570 | ACKR1 | Atypical chemokine receptor 1 | 1 | p<0.0001 | down |
| Q8IZY2 | ABCA7 | ATP binding cassette subfamily A member 7 | 14 | P<0.0001 | down |
| P22234 | PAICS | Isoform 2 of Multifunctional protein ADE2 | 1 | p<0.05 | down |
| P11279 | LAMP1 | Lysosome-associated membrane glycoprotein 1 | 2 | p<0.05 | down |
| O95747 | OXSR1 | Serine/threonine-protein kinase OSR1 | 3 | p<0.05 | down |
| P11498 | PC | Pyruvate carboxylase | 17 | p<0.05 | down |
| P08575 | PTPRC | Receptor-type tyrosine-protein phosphatase C | 7 | p<0.05 | down |
| P62333 | PSMC6 | 26S protease regulatory subunit 10B | 1 | p<0.05 | down |
| P05026 | ATP1B1 | Sodium/potassium-transporting ATPase subunit beta-1 | 2 | p<0.05 | down |
| P10321 | HLA-C | HLA class I histocompatibility antigen, Cw-17 alpha | 1 | p<0.05 | down |
| P07355 | ANXA2 | Isoform 2 of Annexin A2 | 1 | p<0.05 | down |
| Q12913 | PTPRJ | Receptor-type tyrosine-protein phosphatase eta | 2 | p<0.05 | down |
| A6NFX1 | MFSD2B | Major facilitator superfamily domain-containing protein 2B | 2 | p<0.05 | up |
| Q02094 | RHAG | Rh-associated glycoprotein | 2 | p<0.05 | up |
| P78417 | GSTO1 | Glutathione S-transferase omega-1 | 1 | p<0.05 | up |
| O15439 | ABCC4 | Multidrug resistance-associated protein 4 | 41 | p<0.05 | up |
| P04156 | PRNP | Major prion protein | 4 | p<0.05 | up |
| P0C0L5 | C4B | Complement C4-B (Chido blood group) | 2 | p<0.05 | up |
| P02786 | TFRC | Transferrin receptor protein 1 | 3 | p<0.005 | up |

The table shows all (19) differentially expressed proteins between Duffy-positive and -negative RBCs as quantified by Plasma membrane profiling-based Mass spectrometry. Proteins whose abundance is decreased in Duffy-negative relative to the Duffy-positive RBCs are highlighted in blue, while those with increased abundance are highlighted in light grey. A two-tailed t-test was used to determine the level of significance and estimate *P* values, which were corrected for multiple hypothesis testing using the Benjamini–Hochberg method.

**Table S5: Effect of O blood group on *P. falciparum* invasion of Duffy RBCs.**

| Variable | Estimate | SE | *P value* | Summary |
|---|---|---|---|---|
| ***P. falciparum* (3D7)** | | | | |
| Fy phenotype[Fy(a+b+)] | 1.52 | 4.67 | 0.75 | Not significant |
| Fy phenotype[Fy(a+b-)] | -1.59 | 4.46 | 0.73 | Not significant |
| Fy phenotype[Fy(a-b-)] | -6.55 | 4.09 | 0.14 | Not significant |
| ABO[B] | -0.927 | 4.35 | 0.84 | Not significant |
| ABO[O] | -0.322 | 4.16 | 0.94 | Not significant |
| *Adjusted $R^2$=0.02* | | | | |
| ***P. falciparum* (Dd2)** | | | | |
| Fy phenotype[Fy(a+b+)] | -4.03 | 4.25 | 0.36 | Not significant |
| Fy phenotype[Fy(a+b-)] | -5.11 | 4.06 | 0.24 | Not significant |
| **Fy phenotype[Fy(a-b-)]** | **-10.6** | **3.72** | **0.02** | **Significant** |
| ABO[B] | -4.13 | 3.95 | 0.32 | Not significant |
| ABO[O] | 0.37 | 3.79 | 0.92 | Not significant |
| *Adjusted $R^2$=0.33* | | | | |
| ***P. knowlesi*** | | | | |
| Fy phenotype[Fy(a+b+)] | -2.79 | 6 | 0.65 | Not significant |
| Fy phenotype[Fy(a+b-)] | -7.06 | 5.74 | 0.25 | Not significant |
| **Fy phenotype[Fy(a-b-)]** | **-28.3** | **5.26** | **<0.001** | **Significant** |
| ABO[B] | -0.823 | 5.59 | 0.89 | Not significant |
| ABO[O] | 1.46 | 5.35 | 0.79 | Not significant |
| *Adjusted $R^2$=0.70* | | | | |

To rule out any potential confounding effect of the O blood group on the invasion data, we performed a multiple linear regression (MLR) analysis that included the ABO genotypes of the study participants. The analysis showed that the ABO blood groups had no statistically significant impact on the invasion data. However, the Fy(a-b-) phenotype had a minor effect on 3D7 invasion, albeit not statistically significant (*P=0.14*), but had a significant effect on Dd2 invasion (*P=0.02*). As expected, the largest effect of the Duffy-negative phenotype on invasion was noted on the *P. knowlesi* parasites (*P<0.001*). The statistically significant differences are highlighted in bold, and the following blood groups were used as reference: ABO blood group = A, and Duffy phenotype = Fy(a-b+). Adjusted $R^2$ = coefficient of determination, and SE= standard error of the estimate.

### Table S6: FYB$^{ES}$ and sickle cell frequencies in African populations

| Population | Location | HbS frequency | FYB$^{ES}$ frequency | Longitude | Latitude | Subgroup |
|---|---|---|---|---|---|---|
| Mossi | Burkina Faso | 0.034181 | 1 | -1.56159 | 12.23833 | North |
| Bantu | Cameroon | 0.082452 | 0.99498 | 12.35472 | 7.369722 | North |
| Semi-Bantu | Cameroon | 0.063985 | 0.994737 | 12.35472 | 7.369722 | North |
| Fula | Gambia | 0.068069 | 0.998244 | -15.3101 | 13.44318 | North |
| Jola | Gambia | 0.032138 | 0.99927 | -15.3101 | 13.44318 | North |
| Mandinka | Gambia | 0.059532 | 0.998706 | -15.3101 | 13.44318 | North |
| Wollof | Gambia | 0.044461 | 0.999302 | -15.3101 | 13.44318 | North |
| Akans (Ashanti Eastern) | Ghana | 0.037736 | 0.99697 | -1.02319 | 7.946527 | North |
| Kasem | Ghana | 0.022267 | 1 | -1.02319 | 7.946527 | North |
| Nankam | Ghana | 0.029586 | 1 | -1.02319 | 7.946527 | North |
| Northerner | Ghana | 0 | 1 | -1.02319 | 7.946527 | North |
| Bambara | Mali | 0.02451 | 1 | -3.99617 | 17.57069 | North |
| Malinke | Mali | 0.034091 | 1 | -3.99617 | 17.57069 | North |
| Peulh | Mali | 0.04 | 1 | -3.99617 | 17.57069 | North |
| Sarakole | Mali | 0.037037 | 1 | -3.99617 | 17.57069 | North |
| Yoruba | Nigeria | 0.069565 | 1 | 8.675277 | 9.081999 | North |
| Baya-Mandja (Spedini et al. 1981) | Central African Republic | 0.0435 | 0.8528 | 21.36 | 4.77 | SouthCentral |
| Mbugu (Spedini et al. 1981) | Central African Republic | 0.0741 | 0.8841 | 21.36 | 4.77 | SouthCentral |
| Sango (Spedini et al. 1981) | Central African Republic | 0.1227 | 0.8047 | 21.36 | 4.77 | SouthCentral |
| Yakpa (Spedini et al. 1981) | Central African Republic | 0.0694 | 0.8669 | 21.36 | 4.77 | SouthCentral |
| Nyaturu (Godber et al. 1976) | Tanzania | 0.037 | 0.9908 | 35.7 | -4.9 | SouthEast |
| Sandawe (Godber et al. 1976) | Tanzania | 0.0138 | 0.9906 | | | SouthEast |
| Chonyi | Kenya | 0.083167 | 0.998972 | 37.90619 | -0.02356 | SouthEast |
| Giriama | Kenya | 0.035341 | 0.996289 | 37.90619 | -0.02356 | SouthEast |
| Kauma | Kenya | 0.076531 | 1 | 37.90619 | -0.02356 | SouthEast |
| Malawi | Malawi | 0.018003 | 0.994754 | 34.30153 | -13.2543 | SouthEast |
| Mzigua | Tanzania | 0.053571 | 0.997449 | 34.88882 | -6.36903 | SouthEast |
| Wabondei | Tanzania | 0.043956 | 1 | 34.88882 | -6.36903 | SouthEast |
| Wasambaa | Tanzania | 0.038012 | 0.997076 | 34.88882 | -6.36903 | SouthEast |
| Kgalagadi (Jenkins et al. 1987) | Botswana (Kweng District) | 0 | 0.835 | 24.70245 | -23.87 | Southwest |
| Gciriku | Southwest Africa | 0.052 | 0.923 | 19.8 | -17.91 | Southwest |

| | | | | | | |
|---|---|---|---|---|---|---|
| (Nurse and Jenkins 1977) | | | | | | |
| Kwangali (Nurse and Jenkins 1977) | Southwest Africa | 0.034 | 0.905 | 19.8 | -17.91 | Southwest |
| Mbukushu (Nurse and Jenkins 1977) | Southwest Africa | 0 | 0.96 | 19.8 | -17.91 | Southwest |
| Sambyu (Nurse and Jenkins 1977) | Southwest Africa | 0.082 | 0.93 | 19.8 | -17.91 | Southwest |
| Kwengo (Nurse and Jenkins 1977) | Southwest Africa | 0 | 0.885 | 19.8 | -17.91 | Southwest |

Sources are anthropological studies, as indicated in the references, and MalariaGEN SNP data (rs334 in beta globin and rs2814778 in DARC (Malaria Genomic Epidemiology Network et al. 2019); for simplicity, we assume that all instances of rs2814778 "G" represent $FYB^{ES}$, which is a reasonable assumption for African populations). For study (Nurse and Jenkins 1977), the sickle cell allele frequency calculation assumes the number of sickle cell individuals reported is the number of heterozygotes (we believe there to have been a typo in table 9 of study (Nurse and Jenkins 1977) wherein the reported sickle cell allele frequencies of the Sambyu and Gciriku populations are incorrect and we have calculated our own allele frequency values based on the data in the rest of table 9 of that paper).

### Table S7: Parameters of the evolutionary-epidemiological model

| Parameter | Definition | Value used | Notes |
|---|---|---|---|
| $\mu$ | Death rate of hosts from causes other than malaria | 1/25 | mean lifespan of a host = 25 years (intended to be plausible for the period of history over which the evolution of Duffy negativity is likely to have occurred) |
| $\sigma$ | Recovery rate from malaria infection | 2 | mean duration of infection = 6 months (mean durations of infection assumed for *P. falciparum* malaria in epidemiological models range from 20 to 200 days (Mandal et al. 2011); 6 months is in keeping with that assumed by (Filipe et al. 2007), based on 20th century malaria therapy studies. |
| β | Transmission parameter, related to the basic reproduction number ($R_0$) in this model as follows: $R_0 = \frac{\beta}{\sigma+\mu+\alpha_1+\alpha_2}$ | Values between 3 and 11 were tested. | β values were chosen such that $R_0$ takes values between 1.7 and 5. |
| $\alpha_1$ | Severe malaria mortality rate (applies to virulent infections, before immunity to virulence is gained) | 0.01 | For the values of θ, σ, and *r* in this table, if $R_0$ =1.7, these combined mortality rates mean malaria deaths each year are equal to 4.2% of the births. If $R_0$=3, these combined mortality rates mean malaria deaths each year are equal to 4.93% of the births. If $R_0$=5, 5.19% of the births; if $R_0$=10, 5.25% of the births each year. This non-linear relationship between $R_0$ and the cost of malaria to the population occurs because adaptive immunity to the worst virulent effects of malaria is possible within the model, and always gained after the same average number of infections. |
| $\alpha_2$ | Baseline malaria mortality rate (applies to all malaria infections, regardless of immunity to virulence) | 0.00001 | |
| $\psi$ | Reproductive cost to the host of severe infection with malaria | 1 | Severe infection is associated with a 100% probability of reproductive failure (i.e. severe infection whilst pregnant would lead to loss of pregnancy or stillbirth). This figure looks high, but as discussed in (Penman and Gandon 2020), this figure is supposed to capture the reproductive costs of experiencing *P. falciparum* malaria whilst pregnant without any prior adaptive immunity, which is known to be very dangerous. If transmission is high, individuals gain virulence immunity long before reaching reproductive maturity, and do not experience this cost. |
| *g* | Rate at which hosts become reproductively mature | 1/15 | Mean time to reach reproductive maturity = 15 years. |
| *K* | Carrying capacity of population | 10000 | |

| | | | |
|---|---|---|---|
| ***r*** | Fecundity parameter, related to the birth rate as defined in equation 10. | 0.2 | |
| **θ** | Probability of a host gaining immunity to the virulent effects of malaria upon recovery from infection. | 1/15 | This assumes that the average number of infections before gaining immunity to virulence is 15. It is well established that individuals do gain immunity to severe malaria after repeated infection, but the number of infections that is necessary to achieve this is unknown. Figure S9 presents a sensitivity analysis of the model in which we show that the same broad pattern emerges with different values of $\theta$. |
| $p_i$ | Proportion of infections blocked for genotype *i* | Always 0 for the wild type genotype, values for $FYB^{ES}$ homozygotes and heterozygotes based on experimental results from this paper. | |
| $q_i$ | Protection against severe (virulent) malaria mortality enjoyed by genotype *i*. | Always 0 for the wild type genotype, no protection associated with $FYB^{ES}$, but sickle cell heterozygotes are 91% protected against death from severe malaria, based on (Taylor et al. 2012). | |
| $w_i$ | Protection against baseline malaria mortality enjoyed by genotype *i*. | Always 0 for the wild type genotype, no protection associated with $FYB^{ES}$, but sickle cell heterozygotes are 31% protected against baseline malaria mortality, based on (Taylor et al. 2012). | |
